# Genetic drug target validation using Mendelian randomization

**DOI:** 10.1101/781039

**Authors:** A F Schmidt, C Finan, M Gordillo-Marañón, F W Asselbergs, D F Freitag, R S Patel, B Tyl, S Chopade, R Faraway, M Zwierzyna, A D Hingorani

## Abstract

Mendelian randomisation analysis has emerged as an important tool to elucidate the causal relevance of a range of environmental and biological risk factors for human disease. However, inference on cause is undermined if the genetic variants used to instrument a risk factor of interest also associate with other traits that open alternative pathways to the disease (horizontal pleiotropy). We show how the ‘no horizontal pleiotropy assumption’ in MR analysis is strengthened when proteins are the risk factors of interest. Proteins are the proximal effectors of biological processes encoded in the genome, and are becoming assayable on an-omics scale. Moreover, proteins are the targets of most medicines, so Mendelian randomization (MR) studies of drug targets are becoming a fundamental tool in drug development. To enable such studies we introduce a formal mathematical framework that contrasts MR analysis of proteins with that of risk factors located more distally in the causal chain from gene to disease. Finally, we illustrate key model decisions and introduce an analytical framework for maximizing power and elucidating the robustness of drug target MR analyses.

## Introduction

Mendelian randomization (MR) studies estimate the causal relationship of a risk factor of biomedical interest to disease outcomes using genetic variants as instruments to index the risk factor^1^. The naturally randomised allocation of genetic variation at conception reduces the potential for confounding, which compromises causal inference drawn from the directly observed association between risk factor and disease^2^.

Risk factors of biomedical interest (some of which are amenable to modification by drugs or behaviour change) can be both exogenous and endogenous, encompassing health-related behaviours (e.g. smoking and alcohol consumption^3^), complex biological traits (e.g. blood pressure and body mass index^4^) or, circulating constituents of the blood (e.g. complex analytes such as lipoproteins, metabolites such as uric acid^5^, or proteins such as interleukin-6^6,7^, hundreds or thousands of which can now be assayed on high throughput platforms, e.g. from Somalogic and O-Link^8^). Interest has also emerged in tissue-level mRNA expression as an exposure of interest^9^.

Four advances have fuelled an explosion in MR studies. First, genome-wide association studies (GWAS) have provided a rich source of genetic instruments^10^. Second, access to summary level genome wide association data has been made possible through the provision of public data repositories^11^. Third, methods have been developed to execute two-sample MR analysis (risk factor vs disease outcome) based on summary level genetic associations, obviating the need to share potentially sensitive participant information^12^. Fourth, bioinformatics tools have been developed that allow efficient exploitation of such resources. For example, MR-base^13^ is an online platform for MR analyses that links summary level genetic estimates with a number of analytic tools. Additional resources that can be utilised include GCTA, PrediXcan^14^ and MetaXcan^15^.

Most prior MR analyses utilise an approach whereby multiple SNPs identified from GWAS are used as instruments to increase power. SNPs are drawn from throughout the genome, often with a single variant selected per locus^16^ ensuring instruments are independent (i.e. in linkage equilibrium); preventing erroneously inflated statistical significance. This standard approach has often been applied regardless of the position of the exposure of interest in the biological pathway connecting genetic variation to disease risk. For example, an MR analysis investigating the causal relevance of C-reactive protein in a range of disease end-points^17^, and another between educational attainment on cardiovascular disease risk were conducted using broadly similar methodology and assumptions^18^.

However, there are reasons for thinking that MR analysis of a protein risk factor should be considered as a distinct category of MR analysis. First, an analysis of this type induces a natural dichotomy in the genetic instruments that might be used: those that are located in and around the encoding gene (‘*cis*-MR’) vs those located elsewhere in the genome (*trans*-MR) ^19^. Second, aside from mRNA expression, differences in protein expression or function are the most proximal consequence of natural genetic variation. This has two consequences: frequently, variants located in and around the encoding gene can be identified with a very substantial effect on protein expression in comparison to other traits; moreover such instruments may also be less to prone to violating the ‘no horizontal pleiotropy’ assumption’ than variants located elsewhere in the genome (discussed below and ref ^19^). Lastly, in the case of MR analysis of proteins, Crick’s ‘Central Dogma’^20^ imposes an order on the direction of information flow from gene to mRNA to encoded protein, which does not extend beyond this to other biological traits that lie more distally in the causal chain that connects genetic variation to disease risk. Thus, from an MR perspective, proteins are in a privileged position compared to other categories of risk factor.

Understanding which proteins influence which diseases is a fundamental problem in biomedical science since proteins are the major biological effector molecules. Proteins are also the targets of most medicines, so interest has emerged in the use of MR approaches to identify and validate drug targets ^21–24^. Moreover, recent technological developments enable measurement of hundreds or thousands of proteins on an –omics scale in a single biological sample^8^. This opens up the possibility of scaled *cis*-MR analysis of thousands of proteins against hundreds of diseases to inform understanding of their causes and improve drug development yield.

The key to realising this potential is the development of a robust conceptual and mathematical framework for *cis*-MR analysis of proteins. Since *cis*-MR analysis restricts selection of genetic instruments to those located in, or in the vicinity of the encoding gene, new questions emerge as how to optimise the selection of such variants. These include how best to select and define the loci of interest, the physical distance around each gene from which instruments might be drawn; how to select genetic variants as instruments with options including “no selection”, “selection by strength of association”, or “according to functional annotation. Regulatory, non-coding variants act through the level of the encoded protein which is what high-throughput assays detect. Coding-variants might influence protein activity but may also alter the detected rather than actual protein level by protein epitope changes, resulting in a technical artefact. Further questions include whether to weight such instruments in an MR analysis by the level of protein *expression* or *activity*, where the relevant assays are available; or, where they are not, by the level of mRNA expression (and, if so, in which tissue), or by some downstream consequence of protein action, e.g. differences in the level of a metabolite known to be influenced by the protein.

We therefore develop a mathematical framework for *cis*-MR analysis for causal understanding and drug development and investigate the influence of alternative strategies for the selection of genetic instruments and the choice of analytical approach best suited to this task; agnostic of the type of estimation methods.

To help validate our findings, we select examples where the effect of a drug has already been reliably quantified on the protein of interest; on a widely measured downstream mediator of its effect; and on the disease outcome for which the treatment is indicated; and where variants in the gene encoding the drug target have been associated with effects that are consistent with this knowledge. Four genes that fulfil these criteria are *HMGCR*, *PCSK9*, *NPC1L1* and *CETP* that encode the targets of licensed or clinical phase drugs with known effects on lipids and coronary heart disease risk.

## A mathematical framework for *cis*-MR analysis

MR studies determine the causal effect of a risk factor on a disease using instrumental variable (IV) methods^25^, leveraging two estimates: the genetic association with the risk factor (exposure) and the genetic association with the disease (outcome). For the effect estimate in MR to equate to a causal estimate the following critical assumptions should hold: (i) the genetic instrument is (strongly) associated with the exposure, (ii) the genetic instrument is independent of observed and unobserved confounders of the exposure-outcome association (which is secure because genetic variants are fixed and allocated at random), and (iii) conditional on the exposure and confounders, the genetic instrument is independent of the outcome (i.e. there is no instrument – outcome effect other than through the exposure of interest – the “no horizontal pleiotropy” assumption).

The no horizontal pleiotropy assumption is violated when there are additional pathways by which the instrument may be related to the disease, sidestepping the exposure of interest. This could occur, for example if a genetic variant is in linkage disequilibrium (LD) with another variant that influences disease through a pathway distinct from the exposure, or if a genetic instrument also influences disease risk through another risk factor, located proximal to the risk factor of interest in the causal chain from gene to disease. In contrast, the association of a genetic instrument with exposures that lie in the causal chain distal to the exposure of interest (vertical pleiotropy^26^) does not violate the assumptions underpinning MR analysis. In the context of MR analysis of proteins, vertical and horizontal pleiotropy correspond to ‘pre-’ and ‘post’-translational effects respectively.

To address the possibility of horizontal pleiotropy in MR analyses of exposures *other than proteins*, it has been common to select as instruments independently inherited SNPs identified by GWAS from multiple locations across the genome. In doing so, the average horizontal pleiotropy may reduce to zero (so-called *balanced* pleiotropy). Where this is not the case, an estimator such as MR-Egger^27^ can recover an estimate of the causal effect in the presence of horizontal pleiotropy contingent on the Instrument **S**trength is **I**ndependent of the **D**irect **E**ffect (INSIDE) assumption^27^; i.e., that the strength of the genetic association with the risk factor does not determine the magnitude of horizontal pleiotropy..

Figure 1 adapts the typical depiction of an MR analysis to illustrate these considerations. A genetic variant (***G***) has the potential to influence risk of disease (***D***) directly (*ϕ*_***G***_) or through its effect (*δ̃*) on a protein (***P***) which exerts its action through a downstream biomarker (***X***), which in turn influences disease risk. The relevant genetic associations can be resolved as follows:

1. The genetic effect on the protein *δ̃*.
2. The genetic effect on a downstream complex biomarker *δ̃μ*.
3. The genetic effect on disease *ϕ*_***G***_ + *δ̃*(*ϕ*_***P***_ + *μθ*). which comprises:

a. A *direct effect* of the variant on disease *ϕ*_***G***_.
b. An *indirect effect*: *δ̃*(*ϕ*_***P***_ + *μθ*), which is a function of the genetic effect on a protein *δ̃*, the direct effect of the protein on disease *ϕ*_***P***_, the effect of the protein on a biomarker *μ*, and the biomarker effect on disease *θ*.

**Figure 1.**
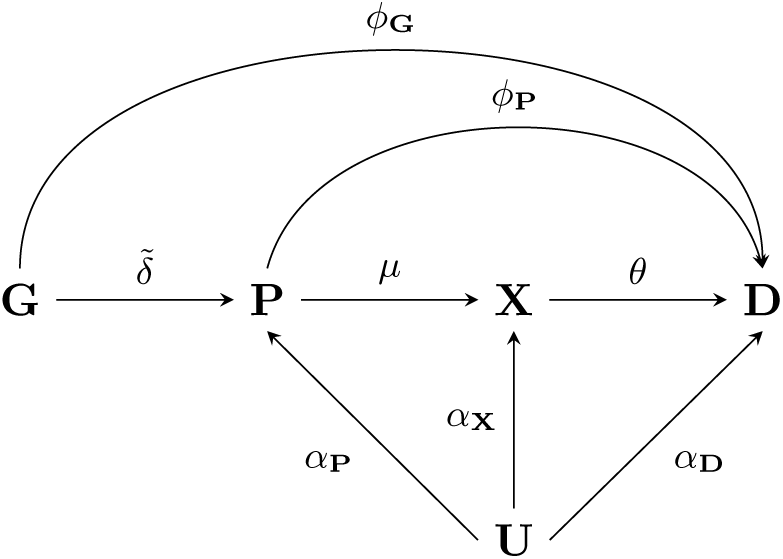
Directed acyclic graphs of potential Mendelian randomization pathways. n.b. nodes are presented in bold face, with **G** representing a genetic variant, **P** a protein drug target, **X** a biomarker, **D** the outcome, and **U** (potentially unmeasured) common causes of both **P**, **X**, **D**. Labelled paths represent the (causal) effects between nodes.

Depending on the risk factor of interest, an MR analysis constitutes a simple quotient of the genetic effect on disease by the genetic effect on the risk factor. For example, if we are interested in the causal effect the biomarker ***X*** on disease, i.e. *θ*, we use the following ratio.

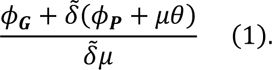

For expression (1) to equate to the causal effect on disease we need to additionally assume that there is no horizontal pleiotropy, in other words *ϕ*_***G***_ = *ϕ*_***P***_ = 0, which reduces the expression to:

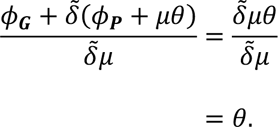

In contrast, if we are interested in the causal effect of the protein ***P*** on disease ***D***, we want to obtain an estimate of *ω*, where *ω* = *ϕ*_***P***_ + *μθ*, we assume *ϕ*_***G***_ = 0 as before, and use the ratio:

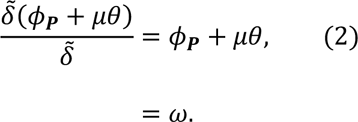

Critically, where the causal effect of the protein is the parameter of interest, we only need to assume that there is no direct effect of the genetic variant on disease, i.e. *ϕ*_***G***_ = 0, and the protein can have any mixture of direct (*ϕ*_***P***_), and indirect (*μθ*) effects. For this reason, MR analysis of protein-disease relationships is less prone to violation of the ‘no horizontal pleiotropy’ assumption than MR analysis of downstream exposures.

### Alternative exposures in cis-MR analysis of proteins

It is important to note that a protein can remain the inferential target in an MR analysis even if it is not measured directly. For example, in cardiovascular disease large sample size GWAS are available on lipids which are often intermediate biomarkers, positioned downstream between the drug target ***P*** and disease, ***D***. In our recent drug target MR analysis of PCSK9 ^22,23^ we used instruments selected from the encoding locus and divided the variant to coronary heart disease (CHD) estimates, not by the effect on PCSK9 level (which was unavailable), but by LDL-C, a variable known to be altered by perturbation of the PCSK9 protein. Thus, using the same notation as above, and assuming *ϕ*_***G***_ = 0:

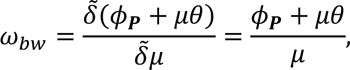

with *bw* indicating ‘biomarker weighted’. Clearly because the denominator contains *δ̃μ*, instead of *δ̃*, *ω_bw_* does not equal *ω*, however *ω_bw_* may still provide a valid null-hypothesis test of *ω* = 0, because *ω_bw_* ≠ 0 implies *ω* ≠ 0, under the assumption of *ϕ*_***G***_ = 0.

In the absence of available measures of the protein of interest, a similar argument can be made for using mRNA expression (this time as an upstream variable) that proxies the effect of genetic variation on the level of the encoded protein (see Appendix Figure 1):

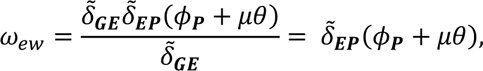

Here the weighting is done by the association with of mRNA expression, and the *δ̃* effect has been decomposed into the variant effect on expression *δ̃*_***GE***_ and the expression effect on protein level *δ̃*_***EP***_ Similar as for *ω_bw_*, the expression weighted (“*ew*”) drug target effect provides a valid test of *ω* = 0 conditional on the absence of any horizontal pleiotropy predicting the protein effect; that is, a necessary assumption *ϕ*_***G***_ = *ϕ*_***E***_ = 0 (with index ***E*** for expression). It should be noted that mRNA expression level (eQTL) is tissue-specific, and utilizing eQTLs for drug target MRs necessitates a decision on the tissue(s) relevant for (de novo) drug development. We return to these issues later in the manuscript.

Having provided the mathematical framework, we now address issues pertaining to the selection of instruments for *cis*-MR analysis of proteins. Specifically, we address locus selection (all protein-coding genes; genes only encoding druggable proteins); locus size; instrument selection (one versus many; selection based on LD or functionality); and potential influence of enhancer variants. Also, we illustrate the influence, and limitations, of weighting genetic instruments by mRNA expression vs. protein expression vs. level of some downstream biomarker, by starting with the current *modus operandi* in MR which is to weight instruments by a downstream biomarker exposure (which is often more widely measured than the protein of interest). Throughout, we draw from current paradigms of MR analysis but critically extend and evaluate modelling choices to derive strategies most relevant for *cis*-MR.

## Design, conduct and interpretation of cis-MR analysis of proteins encoding drug targets

### Selection of loci encoding proteins

Unlike MR analysis of non-protein traits, where it has become common to select instruments from throughout the genome, *cis*-MR analysis necessitates the selection of genetic variants from within or in the vicinity of a protein coding gene. The Ensembl 97 GRCh38 human genome assembly contains an estimated 20,454 protein coding genes, encoding an estimated 24,700 protein coding transcripts (merged Ensembl/Havana annotation). Of these transcripts, 21,869 have a support level of ≤ 2, meaning that there is some level of experimental support for the presence of these transcripts. In addition, UniProt (version 2019_06, combining SwissProt and TrEMBL) reports 20,416 high quality manually annotated proteins.

### Selection of loci encoding druggable genes

Not all encoded proteins are amenable to pharmacological action by small molecule drugs, or peptide and monoclonal antibody therapeutics, which currently account for the majority of medicines. *cis*-MR for drug target validation requires the selection of genes encoding druggable proteins. Progressive efforts to delineate the druggable genome^28,29^ (available through the DGI database (DGIdb^30^), have culminated in the latest iteration containing 4,479 genes^31^ encompassing targets of existing therapeutics, potentially druggable close orthologues and targets accessible by monoclonal antibodies. Of these, ChEMBL v.24 identifies 896 genes as encoding the target components for existing therapeutics, this includes single protein targets, protein complex targets and targets comprising whole protein families. A further 535 genes encode target components of compounds currently in clinical phase testing. Clearly the druggable genome is not static and will be redefined periodically, reflecting changes in drug targeting mechanisms. However, currently, to define the druggable genome is to progressively reduce the high-dimensional search space for genetic instruments from the whole genome to around 20,000 protein coding genes to fewer than 5,000 genes encoding druggable targets. As such, a specific subset of *cis*-MR can inform drug development, which we term “drug target MR”.

### Instrument selection

Drug target MR focuses on a single *gene* known to encode a protein, and *variants* within and around such a gene are used to *characterize* the effect of the drug target on a single or multiple outcome(s). Given the inferential target, it would seem logical to select variants based on the variant to protein level association (*δ̃*). Ideally one would have sound knowledge on the number of causal variants and only select those to minimize bias and maximize precision (power). However, typically this information is unavailable, imposing the need for instrument selection, often using biomarker risk factors as proxies for genetic effects on protein expression.

In such cases, variants are often selected based on 1) a biomarker association (e.g. LDL-C in the case of PCSK9 discussed earlier), 2) predicted functionality; and 3) low linkage disequilibrium (LD). These, often *ad hoc*, selection strategies typically result in the use of a single^24,32,33^ or perhaps a handful of SNPs^22,34,35^ out of a multitude of potential candidate SNPs. Due to a lack of appropriate (pQTL or eQTL) data, it is often unclear how well such a small subset of SNPs characterizes the genetic effect on the drug target (IV assumption i), and how influential selection strategies are on the final MR estimate.

To explore this, we mimicked instrument selection by repeatedly (500 times) sampling four SNPs at random per locus^22,23^ from four known drug target encoding loci *HMGCR* (statins), *NPC1L1* (ezetimibe), *PCSK9* (PCSK9 inhibitors), and *CETP* (CETP inhibitors). (In the next section we consider larger number of variants). These loci contain variants that influence LDL-cholesterol (*HMGCR*, *NPC1L1*, *PCSK9*, *CETP*), with variants at the CETP locus additionally influencing HDL-cholesterol and triglycerides as identified by the Global Lipids Genetics Consortium (GLGC^36^). We then used a generalized least squares (GLS) method^12,37^; to account for pairwise LD between variants at each locus. Variants were extracted from within the gene ±2.5kb, with a minor allele frequency (MAF) above 0.01, and LD less than 0.80 (Appendix Tables 1-5, Appendix Figure 2).

The first and third quartiles (Q) of the CHD odds ratios (OR) per standard deviation (SD) in LDL-C for *HMGCR*, *NPC1L1*, *PCSK9* (or HDL-C in the case of *CETP*) indicated modest variability in the point estimate: (Q1 1.61, Q3 1.78) for *HMGCR*, (Q1 1.42, Q3 1.77) for *PCSK9*, (Q1 1.19, Q3 1.68) for *NPC1L1*, and (Q1 0.87, Q3 0.91) for *CETP*. Between 95-99% of the estimates across all four genes were in the expected direction as inferred from the findings of drugs used in clinical trials to target the corresponding proteins^38–44^.

We further categorised effect estimates based on the EnsEMBL Variant Effect Prediction (VEP) that reports the functional consequence of each variant (Figure 2). We found little to no difference between estimates derived using non-coding variants only and those estimates based on functional variants. The overall stability and agreement between estimates derived with or without functional annotations suggests a strong influence of multivariate LD between the selected and unselected variants in small *cis*-regions (Appendix Figure 2). However, we did observe a large degree of variability in the p-values which is explored in the subsequent section (Appendix Figure3).

**Figure 2.**
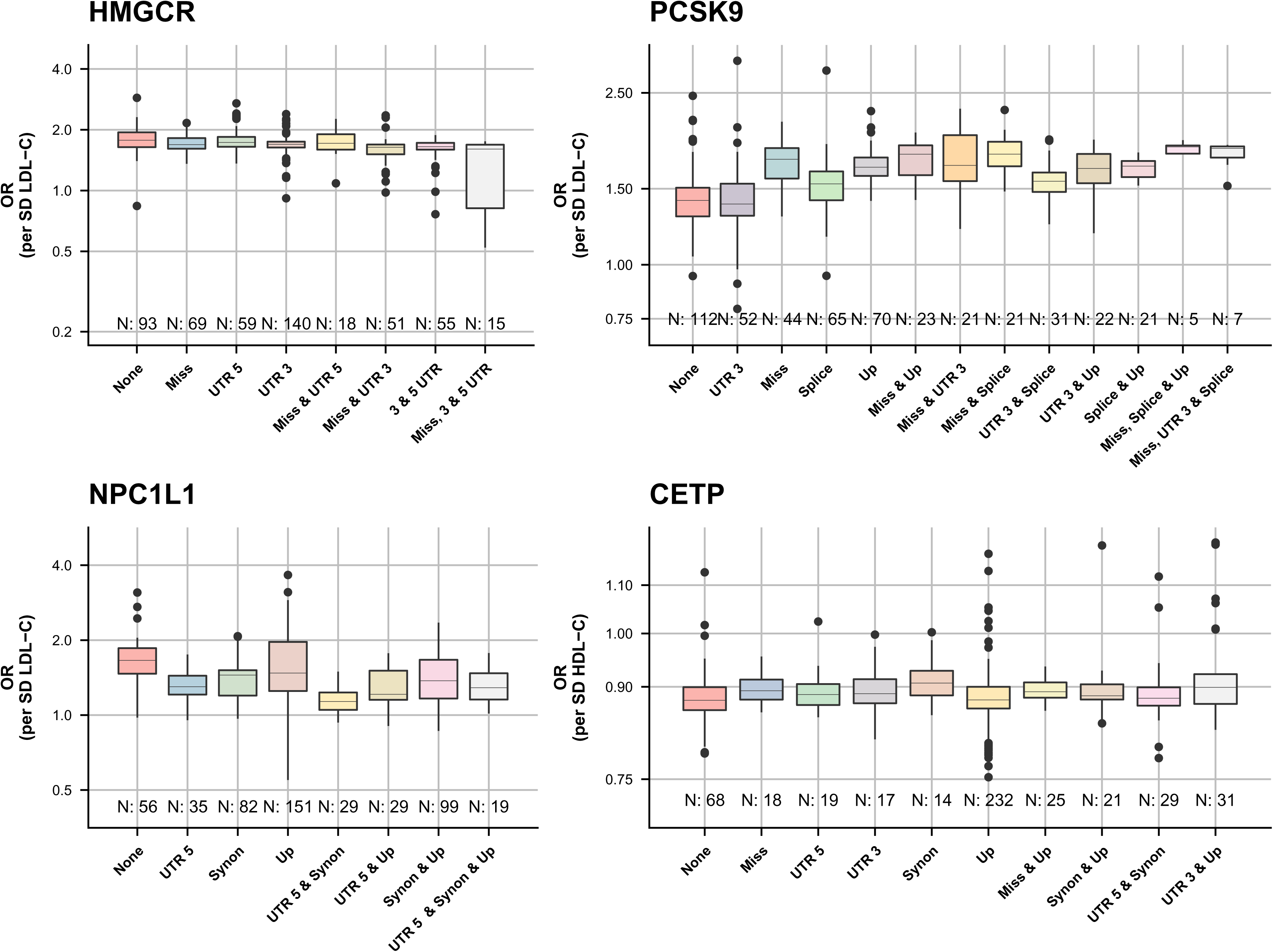
Instrument selection related variation in the point estimates of drug target Mendelian randomization studies on the lipid’s association with CHD. n.b. each estimate is based on randomly (500 iterations) selecting 4 SNPs out of 17 *HMGCR*, 30 *PCSK9*, 21 *NPC1L1*, 36 *CETP* candidate variants. Lipids data was used from the GLGC, and linked to coronary heart disease data from CardiogramPlusC4D. estimates were grouped by the inclusion of instruments with worsted predicted functional or regulatory consequence; categories occurring less than 5 times were removed. Any pairwise LD was accounted for using the 1000 genomes “EUR” reference panel and a generalized least squares method^12^.

### Taking advantage of linkage disequilibrium within the region

Given the observed influence of LD it seems desirable to leverage this in drug target MR. For example, after defining a *cis*-genetic region (discussed further below) one can LD-clump highly correlated variants that might destabilize a statistical model (multicollinearity), and actively model the remaining pairwise LD using an LD-reference panel to maximize power and decrease variability. Besides increasing power and robustness, this strategy also introduces some further complexities e.g. the choice of LD threshold, and the ramification of the possible inclusion of null-variants (i.e. variants that are not associated with the risk-factor).

The effect of LD thresholds can be readily explored by performing a “grid search”, clumping variants at different R-squared thresholds. From modelling theory, (and empirically: Figure 2) one would expect that when using such a grid search that the point estimate stabilizes early (at low thresholds), while the standard errors decrease further until, at a certain point, multicollinearity results in clear deviations. Such a grid search was implemented in Figure 3, showing clear signs of multicollinearity for the *HMGCR* and *PCSK9* estimates, but less so for *NPC1L1*. While trends observed for *HMGCR* and *PCSK9* are examples of what one would expect on theoretical grounds, this does not occur at the same threshold, and seemingly not at all for the *CETP* locus.

**Figure 3.**
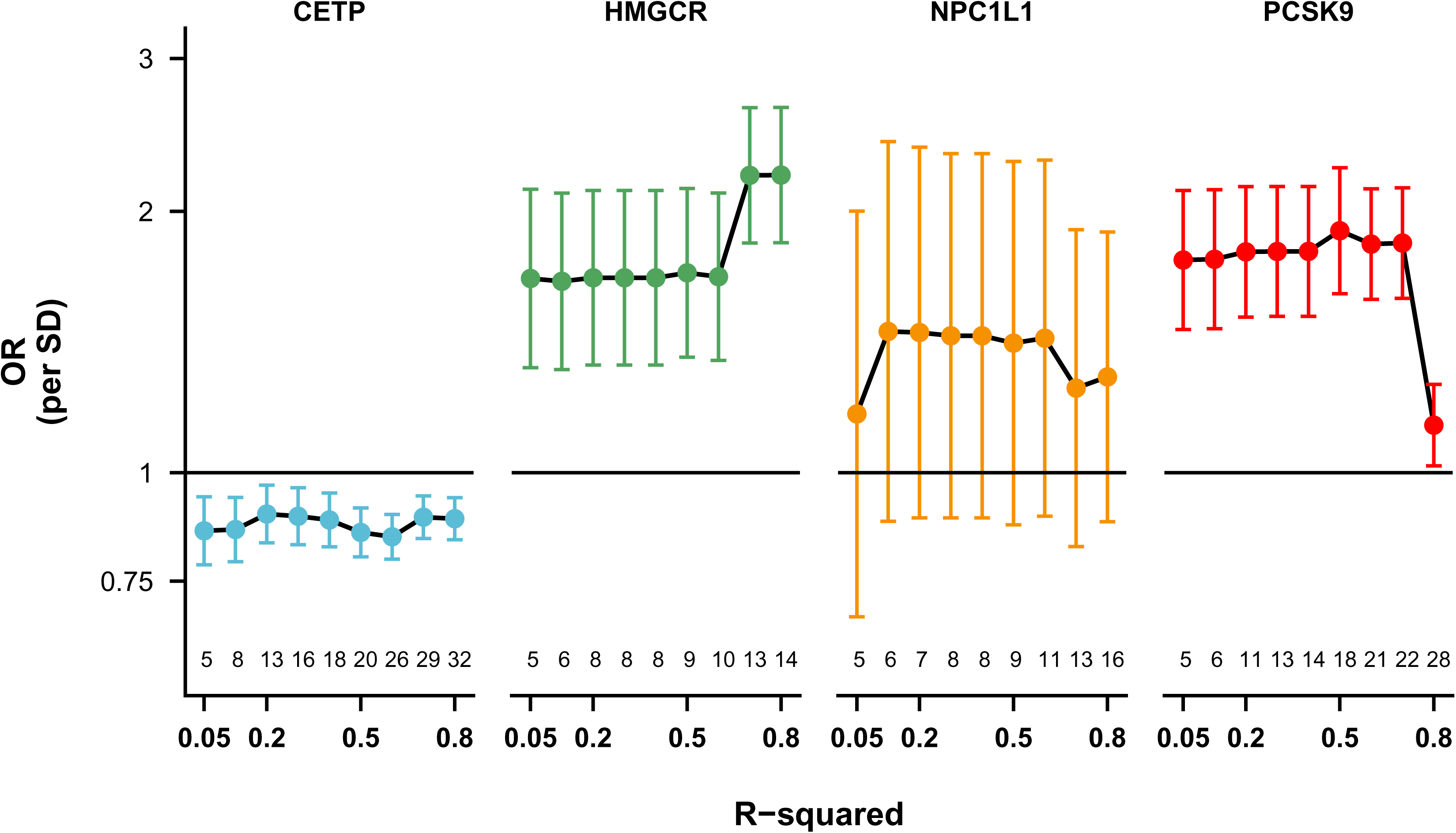
Mendelian randomization estimates of the lipids weighted associations with CHD under increasingly liberal LD-clumping thresholds. n.b. Lipids data was used from the GLGC, and linked to coronary heart disease data from CardiogramPlusC4D. Pairwise LD remaining after LD-clumping was accounted for using the 1000 genomes “EUR” reference panel^56^ and a generalized least squares method^12^. Estimates for *PCSK9*, *HMGCR*, and *NPC1L1* are given per SD in LDL-C, *CETP* estimates per HDL-C reflecting the likely effectiveness pathway to CHD. The number of included variants is depicted and the lower end of the y-axis.

The possible inclusion of “null variants”, that do not affect the intermediate risk factor, is more difficult to prevent. In a very conservative attempt at excluding null-variants researchers often focus on genome-wide significance (e.g., a p-value < 5 × 10^−8^). Dudbridge^45^ and many others have shown that such an approach excludes many useful variants harming power/precision, and lower thresholds (e.g., 10^−5^) often result in greatly improved performance. Clearly such lower threshold could result in the inclusion of (many) null-variants. However, as sample size increases, null-variants will cluster around the origin when regressing the variant-outcome estimates on the variant-exposure estimates, and hence will not affect point estimates of the slope (see Appendix Figure 4). As such, null-variants are not expected to invalidate MR analyses if their inclusion is offset by other variants that are strong predictors of the risk factor; see Appendix Table 11 for a simulation study. Additionally, by employing the two-sample MR^46^ paradigm (using risk factor and outcome estimates from different samples), any possible weak-instrument bias should attenuate results towards the null^47^.

### Linkage disequilibrium modelling compared to selecting functional variants

Based on these considerations, we explored the performance of a very limited instrument selection strategy, geared towards characterizing a *cis*-genetic region encoding a drug target as fully as possible by: 1) considering all variants with limited LD clumping to prevent multicollinearity; 2) modelling LD using external data such as the 1000 genomes reference panel; 3) limited or no p-value thresholding. This strategy (with R^2^ = 0.60) was applied to our four empirical examples and compared to MR estimates at the same locus using only variants with strong evidence of function based on VEP (Figure 4 and Appendix Tables 6-9). We found general agreement between the effect estimates from both analytical approaches, with the GLS estimates having higher precision (1/SE) than those based on functional variants alone. For example, the precision of the two *PCSK9* splice variants used in MR was 5.72 compared to 13.33 for the GLS estimates (incorporating variants selected on the basis of LD structure regardless of function).

**Figure 4.**
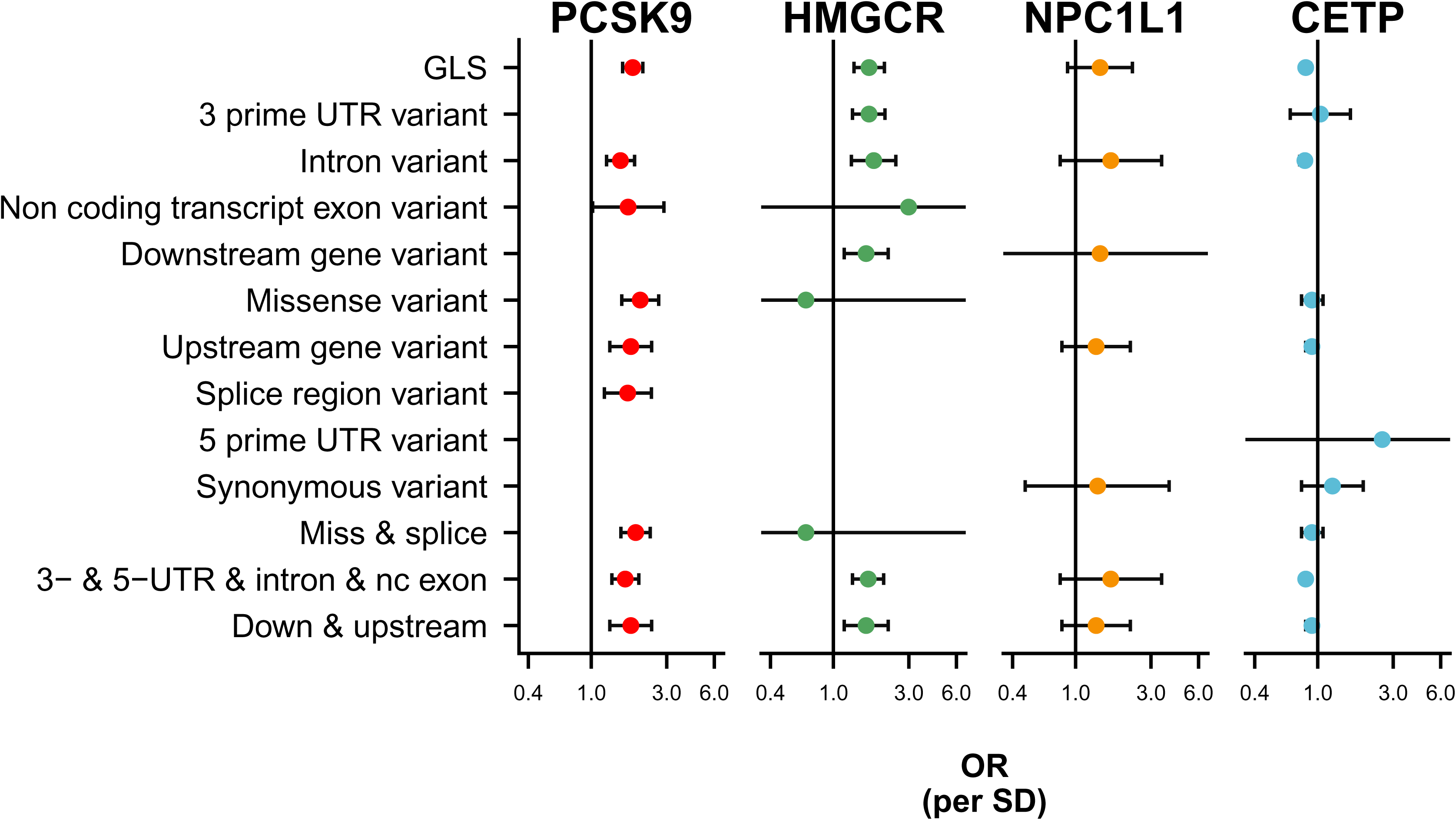
Mendelian randomization estimates of the lipids weighted associations with CHD stratified by functionally of the included variants. n.b. Lipids data was used from the GLGC, and linked to coronary heart disease data from CardiogramPlusC4D. Pairwise LD remaining (after clumping on R-squared of 0.60) was accounted for using the 1000 genomes “EUR” reference panel^56^ and a (GLS) generalized least squares method^12^. Estimates for *PCSK9*, *HMGCR*, and *NPC1L1* are given per SD in LDL-C, *CETP* estimates per HDL-C reflecting the likely effectiveness pathway to CHD.

These results confirm that precision/power is increased by including more correlated variants. To prevent erroneously low p-values in such analyses, we accounted (conditioned) for pairwise LD used the European (EUR) 1000 genomes panel. We further investigated the influence of different 1000G ancestry reference panels on the effect estimates, and found these to be stable for the four examples evaluated (Appendix Figure 5); although significance of the *NPC1L1* was dependent on the panel used. We did find however that the GLS method often failed because of (small) changes in LD resulting in multicollinearity. After inspection this seemed to be related to LD-estimates of low MAF variants varying considerably across ethnicities (Appendix Figure 6); improved behaviour may be expected with either increased sample size (1000G sample size n ∼ 100), or with the removal of low MAF variants (at the risk of losing information).

### Selection of the exposure (risk factor) to be instrumented

The analyses to this point have utilised lipids exposures to index the effect a drug on the corresponding target. With the publication of the INTERVAL study^8^, genetic associations with circulating protein concentration (pQTL data) have been made available for around 3,500 proteins measured using the Somalogic aptamer based chemistry in around 3,000 participants.

This opens up the possibility of using the genetic effect on protein concentration as a more direct proxy of the effect of a drug on its target. Of the four proteins considered here only HMGCR concentrations were available from the INTERVAL study. To supplement this we extracted pQTL estimates from a GWAS of circulating CETP concentration measured by an enzyme-linked immunosorbent assay (ELISA) ^48^ in around 4000 subjects. Initially focussing on the same ±2.5KB region as before, we found the causal estimates for the effect of HMGCR on CHD, using the Somalogic protein expression level to be very imprecise, failing to reject a null-effect (Figure 5); despite the known beneficial effect of HMGCR inhibition by statin drugs on CHD risk. Corresponding estimates using circulating CETP concentration based on an ELISA indicated a causal CHD increasing effect, consistent with the findings of a recent large-scale clinical trial where CETP inhibition reduces CHD risk (Figure 5).

**Figure 5.**
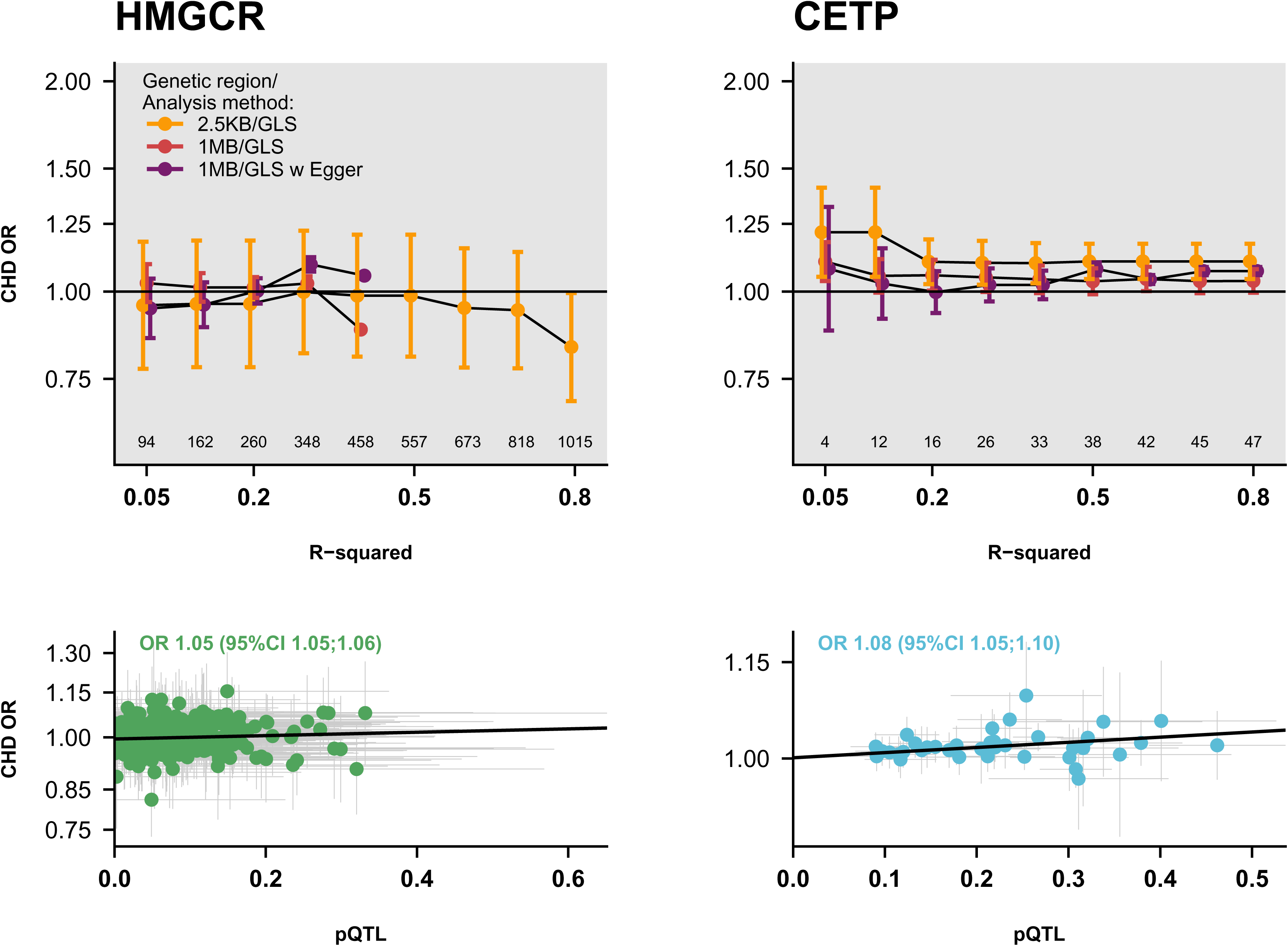
Mendelian randomization estimates of protein level effects on CHD, with a grid of LD threshold. n.b. Pairwise LD was accounted for using the 1000 genomes “EUR” reference panel^56^ and a (GLS) generalized least squares method^12^ with or without Egger correction for possible horizontal pleiotropy. The number of included variants in the 1 mega base flanking region is depicted above the y-axis of the top panels. is depicted and the lower end of the y-axis. The top panel depicts the variant to CHD or protein level effect for clumping threshold 0.3 for HMGCR and 0.5 for CETP, with the GLS Egger slope.

In the biomarker weighted analysis, the size of the genetic flanking region was constrained to prevent erroneously modelling effects from neighbouring genes not encoding the drug target of interest (horizontal pleiotropy). However, pQTL associations provide a direct estimate of the genetic association with the drug-target and hence reduces the need for small flaking regions. We compared findings from the ±2.5KB region, to pQTL MR results using a broader ±1MB flanking region. To further guard against potential horizontal pleiotropy bias (for example through LD) we additionally implemented the Egger adjustment. At intermediate R^2^ values (0.30), HMGCR was significantly associated with an increased CHD risk (with Egger correction). However, with larger values (R^2^ = 0.50), the GLS failed, indicating model instability. Conversely, CETP was (again) robustly causally associated with CHD, with larger R-squared values decreasing variability without any indication of model instability (Figure 5).

We additionally evaluated the performance of MR analysis using mRNA expression level as the exposure variable, assuming a certain proportionality between mRNA and protein expression.

We obtained information on genetic effects on mRNA expression from GTEx^49^ version 7, for all four *cis*-regions, based on post mortem tissues from 449 donors (84% of European descent). Variants were considered that were located in a ±1MB region around each gene. Relative expression levels for each gene differed considerably across tissues and between each locus (Appendix Figure 7). Most strikingly, *HMGCR* was uniformly expressed across tissues, while *CETP* was most expressed in spleen, and *PCKS9* and *NPC1L1* in the liver. In Appendix Figures 8-11 we provide tissue-specific eQTL estimates over a number of genetic regions, showing that *HMGCR* eQTL variants were located throughout the surrounding ±1MB region, and that associations for the other loci were more confined: *CETP* (±10KB), *PCSK9*(±250KB), and *NPC1L1* (±250KB). There was also considerable directional inconsistency in the effects of *cis*-variants on expression across the various tissues, for example *NCP1L1* variants were negatively associated with expression in esophageal mucosa, with the same variants positively associated in aortic artery tissue (Appendix Figure 10).

This directional inconsistency resulted in directionally discordant tissue-specific MR estimates of the same drug-target. For example, PCSK9 mRNA expression in the adrenal gland was associated with an increase in CHD risk: OR 1.0.9 (95%CI 1.02; 1.16), while PCSK9 expression in the uterus was associated with decreased CHD risk: OR 0.92 (95%CI 0.88; 0.97). Selecting variants from a broader ±1MB region universally attenuated effect estimates, with Egger correction (Figure 6) moderately reversing this attenuation. Horizontal pleiotropy did not however fully explain the directional inconsistency in CHD effects across tissues. For example, the causal relationship of HMGCR with CHD weighted based on instrument effects on HMGCR mRNA expression in brain regions (caudate basal ganglia, putamen basal ganglia, and spinal cord cervical c-1 tissues) using Egger correction would be inferred to be protective (*opposite* to that known to be the case from clinical trials).

**Figure 6.**
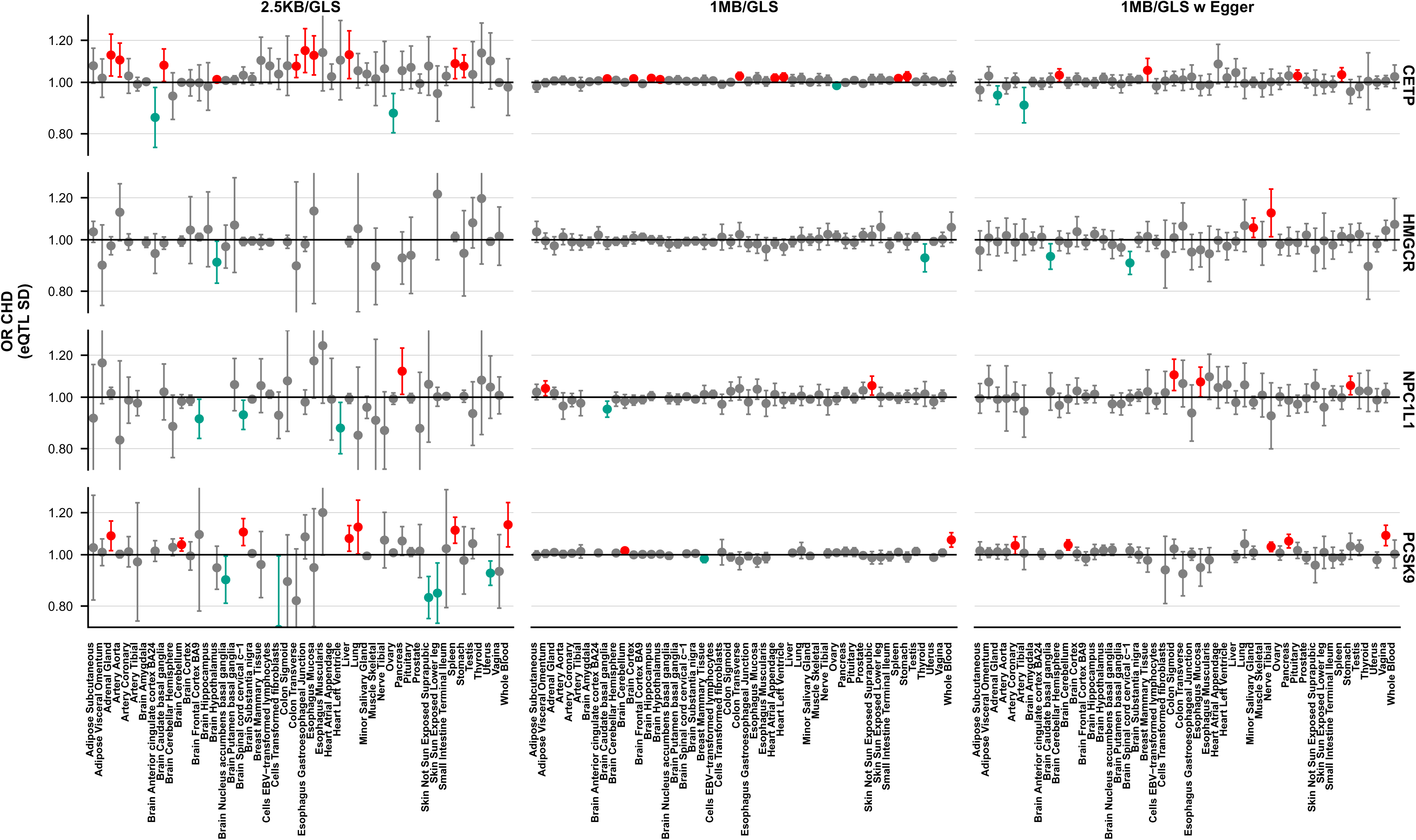
Mendelian randomization estimates utilizing GTEx eQTL weights. Variant were included after clumping on an R-squared threshold of 0.6, with remaining pairwise accounted for using the 1000 genomes “EUR” reference panel^56^ and a generalized least squares method^48^ with, or without accounting for potential pleiotropy using the Egger correction. OR and multiplicative random effects confidence interval per SD chance in expression level are stratified by tissue.

#### Region size

To explore the possible cause of this directional inconsistency we next explored the influence of genetic region by iteratively increasing the flanking region from ±2.5KB to ±1MB, selecting variants from upstream, downstream or in both direction of the gene. With all 4 loci showing similar behaviour, we focus here on the *CETP* region (Figure 7, see appendix Figures 12-14 for the other drug targets) finding that effect direction in any given tissue remained constant across the expanding region. The significance of the association with mRNA expression could either be attenuated when selecting variants from larger regions (e.g. lung tissue), or conversely increase, (e.g., spleen) potentially reflecting different regulatory regions including enhancers for the same gene in different tissues.

**Figure 7.**
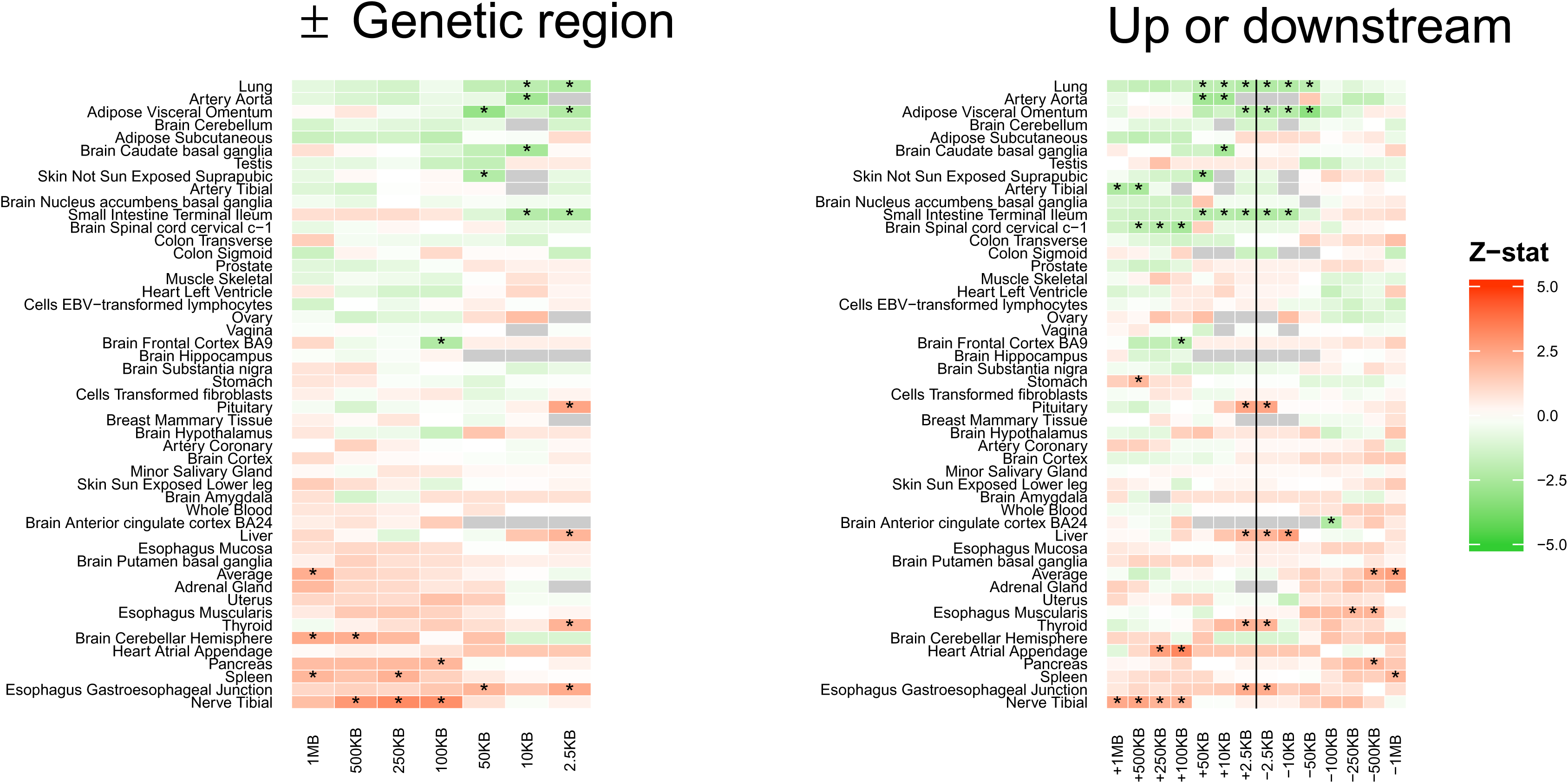
Exploring the influence of the genetic region on the association between CETP expression and CHD. Left hand side selecting from ±the genetic region, right hand side selecting from either the upstream region or the downstream region; colours represent z-statistics direction and stars indicate significant associations at a type 1 error rate of 0.05.

#### Statistical heterogeneity

To further explore the inconsistency in effect direction when weighting MR analysis by mRNA expression in different tissues, we estimated any potential causal relationship between mRNA expression level as the exposure and circulating lipid concentration (rather than CHD risk) as the outcome.

In these analyses, directional inconsistency was also observed (Appendix 15). However, comparing the mRNA expression level effect estimates on lipids, to the expression level estimates on CHD across tissues did not indicate a significant correlation between the two (Appendix 16; correlation estimates between −0.20 and 0.20). As expected both the CHD and lipids estimates showed a large degree of heterogeneity (Appendix Figure 17), indicating either 1) remaining horizontal pleiotropy (despite Egger correction), or 2) true tissue specificity heterogeneity of the expression level effect on CHD. Excluding tissues or variants displaying greater heterogeneity, did not markedly decrease directional inconsistency of estimated causal effects using either lipids or CHD as outcomes (Appendix Figures 18-21).

#### Potential role of enhancer regions

Finally, to determine the influence of enhancer variants, we associated the number of tissue specific enhancers (extracted from HACER^50^) to the MR results from the 4 positive control loci, which did not reveal any significant association (Appendix Figure 22). Additionally, we explored if the number of enhancers could be associated to the direction of effect (encoded as a binary indicator) which did not show a significant association either (Appendix Figure 22). In Figure 8, we associated the tissue-specific heterogeneity statistics of the MR estimates to the distance to the nearest enhancer variants which again did not demonstrate a clear relationship.

**Figure 8.**
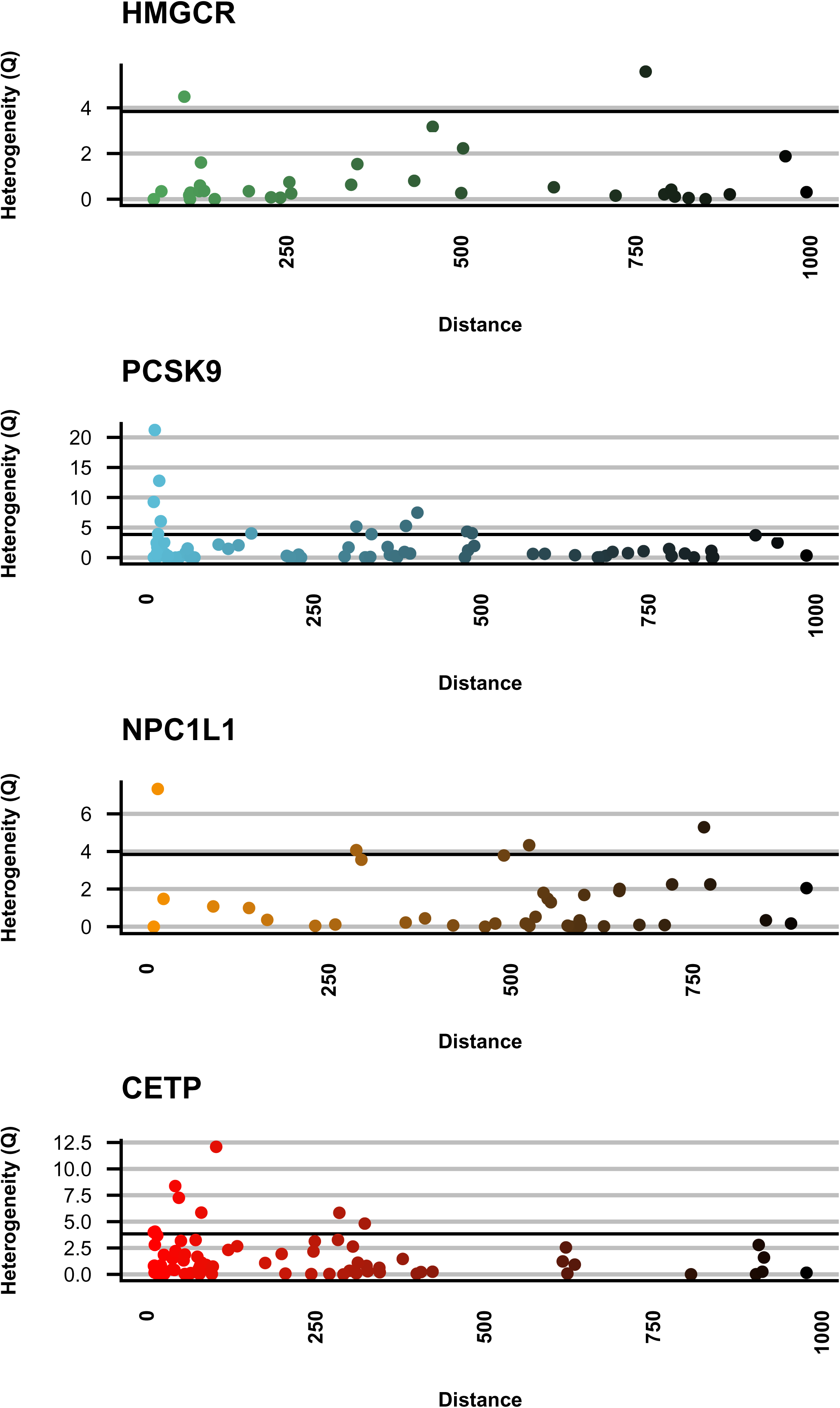
Heterogeneity estimates of the tissue specific MR estimates of the expression level effects on CHD ranked by enhancer distance.

## Discussion

In the current manuscript we have formalised the difference between Mendelian randomization (MR) for biomarker validation and MR for drug target validation *cis*-region encoding a druggable protein. Using algebraic derivations, we show that because drug target MR considers the effects of perturbing a protein drug target on disease, this type of MR may be applied in settings where biomarker MR will be biased through horizontal pleiotropy. We discuss that because drug target MRs can be framed as a *cis*-focus analyses, instrument selection is distinct from that in generic MR, and investigate strategies characterizing the drug target encoding region as whole through linkage disequilibrium (LD), increasing precision and robustness. Simple grid-search algorithms were introduced aiding researcher in optimizing LD-threshold as well as genetic regions, with intuitive sensitivity analyses to determine estimate robustness to the choices of LD reference panel, the presence of functional variants as well as regulatory enhancers, and outliers (using heterogeneity statistics).

Due to our focus on four positive control examples, we were able to perform exhaustive analyses on the robustness of drug target MR findings based on regulatory vs coding variants showing that robust causal inferences could be drawn from regulatory variants despite a widely held view that functional variants should be naturally preferred in (drug target) MR analyses because most drugs affect protein action not level. Instead we showed that modelling an entire *cis* region resulted in the same odds ratio compared to selecting functional variants, with greater precision (increased power) by including larger numbers of (independent) variants. Similarly, we showed that limited influence of LD-reference panels used in LD modelling with non-European ancestry panels resulting in comparable estimates. While promising, these findings should be replicated and above all extended to a larger set of drug targets, for example to analyses outside of the lipids-cardiovascular domain presented.

In the current manuscript we pursued drug target Mendelian randomization by applying a generalized least squares (GLS) solution^51^ to genetic *cis*-regions known to encode protein drug targets. This GLS method is by no means the only relevant estimator function and one can “repurpose” many general MR methods for use in drug target MR. For example, the MR-base platform clumps variants to such a low level (e.g., R-squared of 0.001) that one can apply weighted regression solutions (e.g., IVW), foregoing LD correction^13^. Generalised Summary-data-based Mendelian Randomisation (GSMR)^52^ provides similar LD-modelling MR functionality as the GLS method applied here, which GSMR extents by allowing for automated outlier removal through HEIDI, as well as providing a solid integration with the Genome-wide Complex Trait Analysis (GCTA) suite. Similar outlier removal steps can be readily implemented using the Q-statistic, and standard leverage or Cook’s statistics^53^. Automated outlier removal does however make an implicit assumption that the outlying observations are incorrect, and not the statistical model; which is unlikely to be generally true. Nevertheless, outlier removal is an important step in assessing the robustness of results.

While there are an impressive number of estimator functions, going well beyond the ones just mentioned, there is surprisingly limited advice on what type of instrument should ideally be used for drug target MR (with the exception of Swerdlow *et al.*^19^), with even less attention given to empirically exploring the influence of such advice on MR estimates. This manuscript addresses both issues and introduces a generic framework (Figure 9) for obtaining robust inference of a drug target’s effect on disease, irrespective of the type of MR estimator method preferred. In this framework we suggest that for each exposure outcome pair grid-searches are employed to select the optimal LD threshold and genetic region, while at the same time exploring the robustness of MR estimates to LD-reference panel, the influence of functional and regulatory variants, as well as asses the influence of outlying instruments.

**Figure 9.**
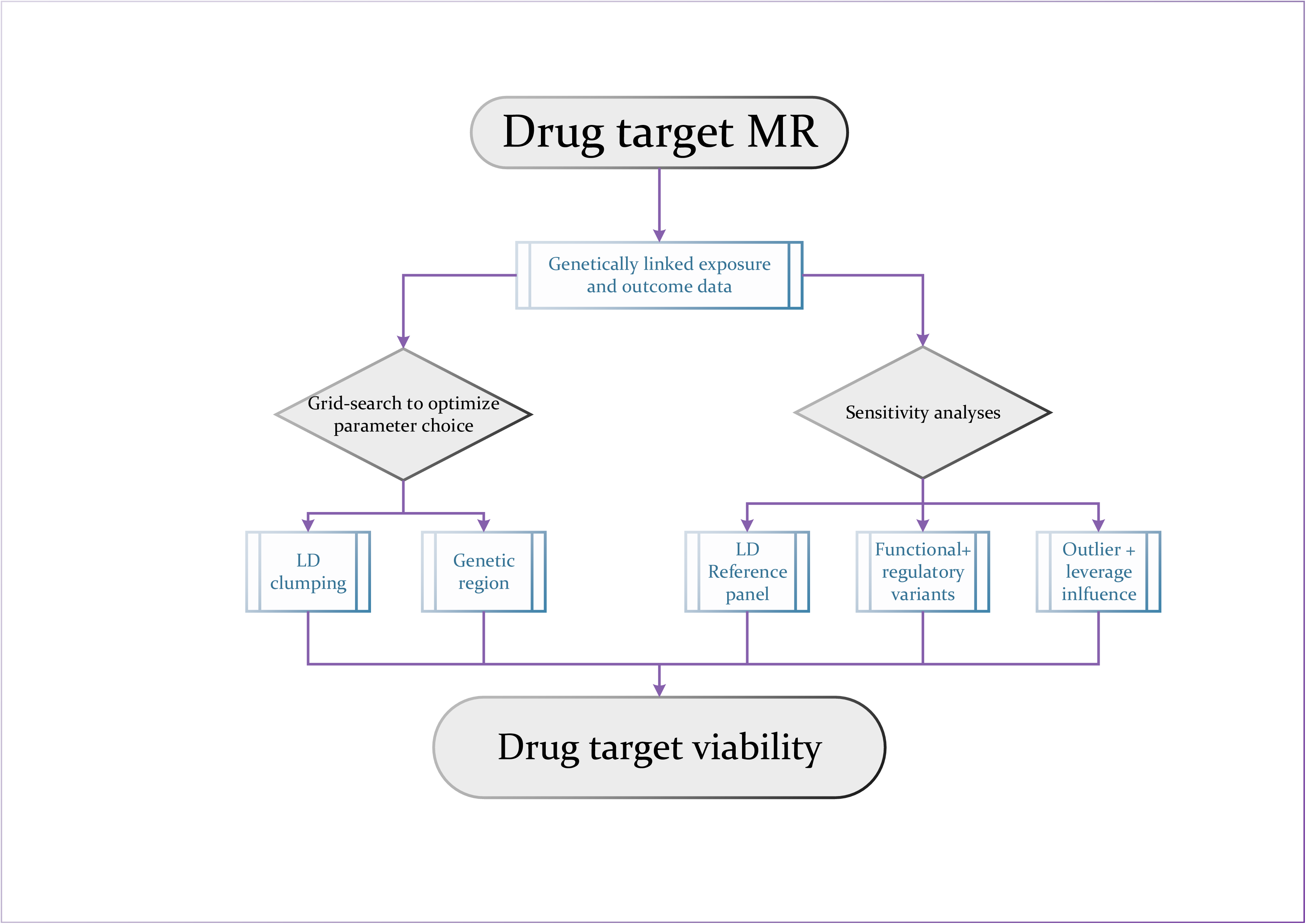
Drug target MR analysis framework.

On theoretical grounds, weighting by protein expression is to be preferred in drug target MR analysis since proteins are the targets of most drugs. Nevertheless, when available, downstream biomarker exposure variables (such as lipids or blood pressure) may provide informative additional exposures, often derived from larger sample sizes than the currently available pQTLs (although this may change in the future). However, many diseases (e.g. neurological disorders) will not have relevant biomarkers and, in the absence of sufficiently large pQTL data, eQTLs provide a seemingly relevant alternative exposure that might closely proxy pQTLs. We have shown however, that eQTL based MR estimates may differ both in magnitude as well as direction across tissues, as demonstrated by exhaustive analyses for the four positive control loci. This tissue-specific heterogeneity was independently reported by the GTEx consortium ^54^ for *PCSK9*. We extend those observations to demonstrate their potential to undermine reliable causal inference when using mRNA expression as a weighting variable in MR analysis. Drug compounds have the potential to affect protein targets in any human tissue in which they are expressed (with the exception of privileged sites such as brain and eyes), however *a priori* it may be unclear which tissue is most relevant for the therapeutic efficacy of a compound, and interpreting the tissue-related heterogeneity may therefore be difficult in pre-clinical settings. It is possible that, despite using methods robust to horizontal pleiotropy, this may have persisted in these analyses, resulting in tissue-specific heterogeneity. Alternatively, the observed heterogeneity may relate to the inclusion of non-European ancestries in the GTEx database^49^, or due to the post-mortem collection of samples^55^. For example, GTEx previously reported that gene expression changed post-mortem in a tissue-specific manner, which they attempted to ameliorate with multiple regression^55^. Without positive control data, however, it is unclear how successful such adjustments have been, and further sensitivity analyses could determine the influence of post-mortem sample interval in MR analysis based on eQTL data.

Throughout this manuscript we used a type I of error rate of 0.05 (or 95% confidence interval) and did not correct for multiple testing. While appropriate multiplicity protection is important, by focussing on four thoroughly studied drug targets (NPC1L1, HMGCR, PCSK9, and CETP) there is an abundance of prior evidence on the expected CHD effect, making *analytical* control of the false positive rate less relevant. In other settings, appropriate control of false discovery rates is clearly essential.it could be argued that applying a genome-wide association p-value threshold (e.g. 5 × 10^−8^) will be needlessly conservative. Instead, of applying the typical GWAS multiplicity correction one could control for the number of druggable proteins (about 5000; resulting in a 1 × 10^−5^ threshold). However, (early) drug development is not performed in isolation, and genetic evidence will be evaluated alongside evidence from cells, tissues, and animal experiments. As such, appropriate false discovery control will depend on the position of drug target MR within this pre-existing evidence framework. For example, p-value threshold of 1 × 10^−5^ might be applied when drug target MR is used as a screening tool, before validating promising leads in further experiments. Positioning drug target MR after successful *in-vivo* experimentation, for example, to check for possible unknown side effects in human subjects, will likely call for a less-stringent multiplicity correction considering themore extensive prior knowledge and the aim of early detection of possible safety concerns.

In conclusion, we expect that combining the discussed concepts with the ever-increasing magnitude of genetic data will move drug target MR from manually curated, often proof of concept like analyses, to more automated and scalable projects able to systematically guide and enrich the entire drug development process.

## Methods

### Data resources

Information on the four positive control loci (*HMGCR, PCSK9, CETP,* and *NPC1L1*) were sourced from the Drug Target Validation Database (DTAdb), developed by Chris Finan^31^. Specifically, for the current analyses we identified variants within a megabase upstream or downstream from each of the four loci. Outcome data were extracted from CARDIOGRAMplusC4D including the genetic association (log odds ratio) with CHD, as well as their standard errors. Exposure data were leveraged from GLGC^36^ (lipids), INTERVAL^8^ (HMGCR protein level), Blauw et al^48^ (CETP protein level), and GTEx^49^ version 7 (expression level).

The 1000 genomes^56^ data were used as a source of LD. Enhancer data were derived from the Human ACtive Enhancer to interpret Regular variants (HACER^50^) resource. All information was curated and normalized to genetic build 37 as described in detail in Finan et al^31^. A ±2.5KB subset of the data is provided in Appendix tables 1-4, with the remainder easily extracted from cited publicly available sources.

### Statistical analysis

Mendelian randomization was conducted using the “Inverse Variance Weighted” (IVW) and “MR-Egger” methods for correlated variants as detailed in Burges et al^37^. Here we note that these methods are specific parametrizations of Generalized Least Squares (GLS) technique and simply refer to IVW as GLS, and MR-Egger as GLS with Egger correction.

In the context of MR, a GLS without an Egger correct, regresses the genetic association with an outcome (CHD in our case) on the genetic association with an exposure (here lipids, protein level, or exposure level), forcing the intercept through zero; reflecting the no-pleiotropy assumption when a zero exposure effect should be matched by a zero outcome effect. Here the slope estimate equates to a causal estimate of the exposure to outcome effect. GLS with Egger correction refers to a similar linear model without forcing the intercept through the origin. Here the intercept estimate reflects the amount of horizontal pleiotropy, while the slope estimates reflects the causal estimate of the exposure on the outcome corrected for (potential) horizontal pleiotropy). Estimates are presented as fixed effects (with a regression standard error of unity), or as random effects (where the regression standard error is equal or larger than 1).

All analyses were conducted using the R programming language^57^, with packages dplyr^58^, ggplot2^59^, gridExtra^60^, openxlsx^61^, and wesanderson^62^. Diagrams were programmed in TikZ^63^, and the appendix written in LaTeX and knitr^64^.

## Acknowledgement

AFS is supported by BHF grant PG/18/5033837 and the UCL BHF Research Accelerator AA/18/6/34223. CF and AFS received additional support from the National Institute for Health Research University College London Hospitals Biomedical Research Centre. ADH is an NIHR Senior Investigator. We further acknowledge support from the the Rosetrees and Stoneygate Trust. FWA is supported by UCL Hospitals NIHR Biomedical Research Centre. RSP is supported by a BHF Fellowship FS/14/76/30933. This work has received support from the EU/EFPIA Innovative Medicines Initiative [2] Joint Undertaking BigData@Heart grant n° 116074.

## Author contributions

AFS, CF, MGM, RF, MZ, and ADH contributed to the idea, and design of the study, AFS, CF, SC, and MGM performed the analyses and drafted the manuscript. CF and SC curated, extracted and normalized these data. AFS and CF drafted the manuscript. MG, RP, DFF, BT, FWA, SC, RF, MZ and ADH provided critical input on the analyses and the drafted manuscript.

## Prior postings and presentations

This study and its results have been submitted to bioRxiv XXX, and an abstract has been submitted to the 2019 Mendelian Randomization conference in Bristol as well as to the EMBL “Target Validation Using Genomics and Informatics” conference.

## Data availability

All data are publicly available, as described in the methods section. The data specifically used for these analyses is available through (URL will be made available after acceptance)

---

**Table 1.** Glossary

| Term | Description |
| --- | --- |
| Confounder | A common cause of both the exposure and outcome. Ignored confounding (measured or unmeasured) biases the exposure to outcome association. |
| Estimand | The parameter to be estimated. For example, the population mean $\mu$ |
| Estimate | The result of an estimator function. For example, the sample estimate $\hat{\mu}$ |
| Estimator | A function that provides an estimate of an estimand. For example, an estimator of the population mean would be $\frac{1}{n} \sum_{i=1}^n x_i$ for $n$ measurements. |
| Exposure | A phenotype which may cause an outcome trait (disease state or risk factor). In the manuscript often refers to mRNA expression, protein, or biomarker levels. |
| Generalised least squares (GLS) | Generalised least squares is an extension of ordinary least squares (linear regression) to account for correlation between variants (due to LD). In the manuscript GLS refers to a linear model regressing the variant to outcome association on the variant to exposure association with the intercept forced through the origin. A GLS with Egger correction (see MR-Egger) refers to a similar linear model with the intercept is a free parameter. |
| Inverse variance weighted (IVW) | The IVW is an IV-estimator assuming an absence of directional pleiotropy. Can be implement using a GLS with the intercept forced through the origin. |
| Mendelian randomization | A study or principle that use genetic instrumental variables, often in a formal instrumental variable analysis. |
| Mendelian randomization for biomarker validation |  |
| Drug target Mendelian randomization |  |
| MR-Egger | MR-egger is an IV-estimator which account for potential directional pleiotropy by allowing the intercept of an (weighted) OLS to differ from zero. |
| Intermediate exposure or intermediate phenotype | See exposure; Intermediate refers to the assumed position of the exposure between the genetic variant and the (outcome) trait. |
| Instrument | Any variable that fulfils the three instrumental variable (IV) assumptions:<br>1. The instrument is strongly associated with exposure<br>2. The instrument is independent of observed and unobserved confounders of the exposure-outcome relation.<br>3. Conditional on exposure and confounders, the instrument is independent of the outcome. |
| Instrumental variable | See instrumental variable |
| Instrumental variable analysis | A method to estimate the causal effect of a potentially confounded exposure-outcome relation. |
| Outcome | A disease state, risk factor, or general phenotype that is thought to be caused by an exposure or genetic variant. |
| P-value as a random variable | A p-value can be interpreted as a random variable, meaning that a single experiment can have a randomly high or low value (centred around an unknown mean, with unknown standard deviation). |
| Pleiotropy | A genetic variant being associated with multiple phenotypes. Vertical pleiotropy refers to variants being associated to traits in the same pathway (or cascade), for example, *PCSK9* → LDL-C → CHD. Horizontal pleiotropy occurs when a genetic variant is associated with the outcome of interest via direct pathway that is not mediated by the exposure; in the previous example this would be *PCSK9* affecting CHD via a non-lipids pathway. |

## References

1. Lawlor, D. A. et al. Mendelian randomization: using genes as instruments for making causal inferences in epidemiology. Stat.Med. 27, 1133–1163 (2008).

2. Hingorani, A. & Humphries, S. Nature’s randomised trials. Lancet 366, 1906–1908 (2005).

3. Holmes, M. V et al. Association between alcohol and cardiovascular disease: Mendelian randomisation analysis based on individual participant data. BMJ 349, g4164 (2014).

4. Holmes, M. V. et al. Causal effects of body mass index on cardiometabolic traits and events: A Mendelian randomization analysis. American Journal of Human Genetics 94, 198–208 (2014).

5. White, J. et al. Plasma urate concentration and risk of coronary heart disease: A Mendelian randomisation analysis. The Lancet Diabetes and Endocrinology 4, 327–336 (2016).

6. Swerdlow, D. I. et al. The interleukin-6 receptor as a target for prevention of coronary heart disease: A mendelian randomisation analysis. The Lancet 379, 1214–1224 (2012).

7. Sarwar, N. & Butterworth, A. S. Interleukin-6 receptor pathways in coronary heart disease: A collaborative meta-analysis of 82 studies. The Lancet 379, 1205–1213 (2012).

8. Sun, B. B. et al. Genomic atlas of the human plasma proteome. Nature (2018). doi:10.1038/s41586-018-0175-2

9. Xu, X. et al. Molecular insights into genome-wide association studies of chronic kidney disease-defining traits. Nature Communications 9, 4800 (2018).

10. Buniello, A. et al. The NHGRI-EBI GWAS Catalog of published genome-wide association studies, targeted arrays and summary statistics 2019. Nucleic Acids Res 47, D1005–D1012 (2019).

11. Kamat, M. A. et al. PhenoScanner V2: an expanded tool for searching human genotype–phenotype associations. Bioinformatics doi:10.1093/bioinformatics/btz469

12. Burgess, S., Dudbridge, F. & Thompson, S. G. Combining information on multiple instrumental variables in Mendelian randomization: Comparison of allele score and summarized data methods. Statistics in Medicine 35, 1880–1906 (2016).

13. Hemani, G. et al. The MR-Base platform supports systematic causal inference across the human phenome. eLife (2018). doi:10.7554/eLife.34408

14. Gamazon, E. R. et al. A gene-based association method for mapping traits using reference transcriptome data. Nature Genetics 47, 1091–1098 (2015).

15. MetaXcan: Summary Statistics Based Gene-Level Association Method Infers Accurate PrediXcan Results.

16. Davies, N. M., Holmes, M. V. & Davey Smith, G. Reading Mendelian randomisation studies: a guide, glossary, and checklist for clinicians. BMJ k601 (2018). doi:10.1136/bmj.k601

17. Ligthart, S. et al. Genome Analyses of >200,000 Individuals Identify 58 Loci for Chronic Inflammation and Highlight Pathways that Link Inflammation and Complex Disorders. The American Journal of Human Genetics 103, 691–706 (2018).

18. Tillmann, T. et al. Education and coronary heart disease: mendelian randomisation study. BMJ 358, j3542 (2017).

19. Swerdlow, D. I. et al. Selecting instruments for Mendelian randomization in the wake of genome-wide association studies. International journal of epidemiology 45, 1600–1616 (2016).

20. Crick, F. Central dogma of molecular biology. Nature 227, 561–563 (1970).

21. Zheng, J. et al. Phenome-wide Mendelian randomization mapping the influence of the plasma proteome on complex diseases. bioRxiv 627398 (2019). doi:10.1101/627398

22. Schmidt, A. F. et al. PCSK9 genetic variants and risk of type 2 diabetes: a mendelian randomisation study. The Lancet Diabetes & Endocrinology 0, 735–742 (2016).

23. Schmidt, A. F., et al. Phenome-wide association analysis of LDL-cholesterol lowering genetic variants in PCSK9. bioRxiv (2018).

24. Swerdlow, D. I. et al. HMG-coenzyme A reductase inhibition, type 2 diabetes, and bodyweight: Evidence from genetic analysis and randomised trials. The Lancet 385, 351–361 (2015).

25. Boef, A. G. C., Dekkers, O. M. & Le Cessie, S. Mendelian randomization studies: A review of the approaches used and the quality of reporting. International Journal of Epidemiology 44, 496–511 (2015).

26. Hemani, G., Bowden, J. & Davey Smith, G. Evaluating the potential role of pleiotropy in Mendelian randomization studies. Hum. Mol. Genet. 27, R195–R208 (2018).

27. Bowden, J., Smith, G. D. & Burgess, S. Mendelian randomization with invalid instruments: Effect estimation and bias detection through Egger regression. International Journal of Epidemiology 44, 512–525 (2015).

28. Hopkins, A. L. & Groom, C. R. The druggable genome. Nature reviews. Drug discovery 1, 727–30 (2002).

29. Russ, A. P. & Lampel, S. The druggable genome: An update. Drug Discovery Today (2005). doi:10.1016/S1359-6446(05)03666-4

30. Cotto, K. C. et al. DGIdb 3.0: a redesign and expansion of the drug-gene interaction database. Nucleic Acids Res. 46, D1068–D1073 (2018).

31. Finan, C. et al. The druggable genome and support for target identification and validation in drug development. Science Translational Medicine 9, (2017).

32. Swerdlow, D. I., Hingorani, A. D., Casas, J. P. & Consortium, I. M. R. The interleukin-6 receptor as a target for prevention of coronary heart disease: a mendelian randomisation analysis. Lancet 379, 1214–1224 (2012).

33. Sofat, R. et al. Separating the mechanism-based and off-target actions of cholesteryl ester transfer protein inhibitors with CETP gene polymorphisms. Circulation 121, 52–62 (2010).

34. Ference, B. A. et al. Variation in PCSK9 and HMGCR and Risk of Cardiovascular Disease and Diabetes. New England Journal of Medicine 375, 2144–2153 (2016).

35. Ference, B. A., Majeed, F., Penumetcha, R., Flack, J. M. & Brook, R. D. Effect of naturally random allocation to lower low-density lipoprotein cholesterol on the risk of coronary heart disease mediated by polymorphisms in NPC1L1, HMGCR, or Both: A 2 by 2 factorial mendelian randomization study. Journal of the American College of Cardiology 65, 1552–1561 (2015).

36. Willer, C. J. et al. Newly identified loci that influence lipid concentrations and risk of coronary artery disease. 40, (2008).

37. Burgess, S., Zuber, V., Valdes-Marquez, E., Sun, B. B. & Hopewell, J. C. Mendelian randomization with fine-mapped genetic data: Choosing from large numbers of correlated instrumental variables. Genetic Epidemiology 41, 714–725 (2017).

38. Schmidt, A. F. A. F. et al. PCSK9 monoclonal antibodies for the primary and secondary prevention of cardiovascular disease. Cochrane Database of Systematic Reviews 2017, (2017).

39. Cholesterol Treatment Trialists’ (CTT) Collaborators, C. T. T. (CTT) et al. The Effects of Lowering LDL Cholesterol with Statin Therapy in People at Low Risk of Vascular Disease: Meta-Analysis of Individual Data from 27 Randomised Trials. Journal of Vascular Surgery 57, 284 (2013).

40. Collins, R. et al. Interpretation of the evidence for the efficacy and safety of statin therapy. The Lancet 388, 2532–2561 (2016).

41. Keene, D., Price, C., Shun-Shin, M. J. & Francis, D. P. Effect on cardiovascular risk of high density lipoprotein targeted drug treatments niacin, fibrates, and CETP inhibitors: meta-analysis of randomised controlled trials including 117,411 patients. BMJ : British Medical Journal 349, g4379- (2014).

42. Bohula, E. A. et al. Prevention of Stroke with the Addition of Ezetimibe to Statin Therapy in Patients with Acute Coronary Syndrome in IMPROVE-IT. Circulation CIRCULATIONAHA.117.029095 (2017). doi:10.1161/CIRCULATIONAHA.117.029095

43. Cannon, C. P. et al. Ezetimibe Added to Statin Therapy after Acute Coronary Syndromes. N Engl J Med 372, 2387–2397 (2015).

44. Schmidt, A. F., Pearce, L. S., Wilkins, J. T., Casas, J. P. & Hingorani, A. D. Cochrane corner: PCSK9 monoclonal antibodies for the primary and secondary prevention of cardiovascular disease. Heart 104, 1053 LP–1055 (2018).

45. Dudbridge, F. Power and Predictive Accuracy of Polygenic Risk Scores. PLoS Genetics 9, (2013).

46. Bowden, J. et al. Assessing the suitability of summary data for two-sample Mendelian randomization analyses using MR-Egger regression: the role of the I2 statistic. International journal of epidemiology 1–14 (2016). doi:10.1093/ije/dyw220

47. Burgess, S. & Thompson, S. G. Avoiding bias from weak instruments in mendelian randomization studies. International Journal of Epidemiology 40, 755–764 (2011).

48. Blauw, L. L., et al. CETP (Cholesteryl Ester Transfer Protein) Concentration: A Genome-Wide Association Study Followed by Mendelian Randomization on Coronary Artery Disease. Circ Genom Precis Med 11, e002034 (2018).

49. Aguet, F. et al. Genetic effects on gene expression across human tissues. Nature (2017). doi:10.1038/nature24277

50. Liu, Y., Sarkar, A., Kheradpour, P., Ernst, J. & Kellis, M. Evidence of reduced recombination rate in human regulatory domains. Genome Biology 18, 193 (2017).

51. Burgess, S., Zuber, V., Valdes-Marquez, E., Sun, B. B. & Hopewell, J. C. Mendelian randomization with fine-mapped genetic data: Choosing from large numbers of correlated instrumental variables. Genetic Epidemiology 41, 714–725 (2017).

52. Zhu, Z. et al. Causal associations between risk factors and common diseases inferred from GWAS summary data. Nat Commun 9, 1–12 (2018).

53. Schmidt, A. F. & Finan, C. Linear regression and the normality assumption. Journal of Clinical Epidemiology 98, 146–151 (2018).

54. Barbeira, A. N. et al. Exploring the phenotypic consequences of tissue specific gene expression variation inferred from GWAS summary statistics. Nature Communications (2018). doi:10.1038/s41467-018-03621-1

55. Ferreira, P. G. et al. The effects of death and post-mortem cold ischemia on human tissue transcriptomes. Nature Communications (2018). doi:10.1038/s41467-017-02772-x

56. The 1000 Genomes Project Consortium. A global reference for human genetic variation. Nature 526, 68–74 (2015).

57. R Core Team. R: A language and environment for statistical computing. (R Foundation for Statistical Computing, 2017).

58. Hadley Wickham, Lionel Henry, Kirill Müller & Romain François. dplyr: A Grammar of Data Manipulation. (2019).

59. Wickham, H. ggplot2: elegant graphics for data analysis. (Springer, 2016).

60. Baptiste Auguie. gridExtra: Miscellaneous Functions for ‘Grid’ Graphics. (2017).

61. Alexander Walker. openxlsx: Read, Write and Edit XLSX Files. (2019).

62. Karthik Ram & Hadley Wickham. wesanderson: A Wes Anderson Palette Generator. (2018).

63. Till Tantau. The TikZ and PGF Packages. (2013).

64. Xie, Y. Dynamic documents with R and Knitr. (CRC Press/Taylor & Francis, 2015).

