## Appendix for "Genetic drug target validation using Mendelian randomization"

**Appendix Table 1:** Genetic associations with pQTL, LDL-C, and CHD, in a $\pm 2.5$KB region around the *HMGCR* locus (clumped on $R^2 < 0.80$). GWAS data was extracted from INTERVAL[1], GLGC[2], and Cardiogram-PlusC4D[3, 4].

| rsID | Mean difference pQTL | SE pQTL | Mean difference per SD LDL | SE per SD LDL | logOR CHD | SE CHD | Allele frequency | Worst consequence |
|---|---|---|---|---|---|---|---|---|
| rs10473972 | 0.0135 | 0.0285 | NA | NA | -0.023 | 0.011 | 0.259 | intron variant |
| rs10474435 | NA | NA | -0.054 | 0.015 | 0.021 | 0.035 | 0.011 | 3 prime UTR variant |
| rs10515198 | NA | NA | -0.060 | 0.006 | -0.039 | 0.014 | 0.100 | intron variant |
| rs112585543 | 0.0991 | 0.1036 | NA | NA | -0.007 | 0.042 | 0.011 | intron variant |
| rs115169875 | 0.1117 | 0.0790 | 0.034 | 0.015 | -0.028 | 0.051 | 0.021 | intron variant |
| rs12916 | NA | NA | -0.073 | 0.004 | -0.039 | 0.009 | 0.413 | 3 prime UTR variant |
| rs138091235 | -0.0448 | 0.1123 | NA | NA | 0.057 | 0.042 | 0.013 | intron variant |
| rs142550951 | -0.0947 | 0.0568 | NA | NA | -0.007 | 0.025 | 0.056 | intron variant |
| rs17238540 | 0.0061 | 0.0820 | 0.024 | 0.016 | 0.026 | 0.039 | 0.017 | non coding transcript exon variant |
| rs17238568 | -0.1047 | 0.0474 | -0.032 | 0.010 | -0.023 | 0.021 | 0.075 | intron variant |
| rs17238596 | 0.0463 | 0.0624 | NA | NA | -0.001 | 0.039 | 0.044 | intron variant |
| rs17244792 | -0.0842 | 0.1009 | NA | NA | -0.035 | 0.056 | 0.018 | intron variant |
| rs17244848 | -0.0484 | 0.0400 | NA | NA | -0.027 | 0.017 | 0.105 | intron variant |
| rs17244939 | 0.1151 | 0.0904 | 0.054 | 0.020 | 0.039 | 0.064 | 0.013 | intron variant |
| rs17244953 | -0.0709 | 0.0633 | 0.005 | 0.013 | -0.018 | 0.031 | 0.036 | 5 prime UTR variant |
| rs17648121 | 0.0994 | 0.0792 | NA | NA | -0.006 | 0.033 | 0.030 | intron variant |
| rs2241402 | -0.0394 | 0.0561 | -0.033 | 0.011 | -0.028 | 0.023 | 0.051 | intron variant |
| rs2303152 | NA | NA | -0.042 | 0.006 | -0.021 | 0.014 | 0.110 | intron variant |

**Appendix Table 1:** Genetic associations with pQTL, LDL-C, and CHD, in a $\pm 2.5$KB region around the *HMGCR* locus (clumped on $R^2 < 0.80$). GWAS data was extracted from INTERVAL[1], GLGC[2], and Cardiogram-PlusC4D[3, 4]. *(continued)*

| rsID | Mean difference pQTL | SE pQTL | Mean difference per SD LDL | SE per SD LDL | logOR CHD | SE CHD | Allele frequency | Worst consequence |
|---|---|---|---|---|---|---|---|---|
| rs3761739 | NA | NA | -0.046 | 0.005 | -0.024 | 0.012 | 0.152 | intron variant |
| rs3846662 | -0.014 | 0.0255 | NA | NA | -0.032 | 0.009 | 0.432 | non coding transcript exon variant |
| rs4629571 | -0.0767 | 0.0402 | NA | NA | -0.010 | 0.016 | 0.111 | upstream gene variant |
| rs4703670 | 0.0314 | 0.0306 | -0.062 | 0.004 | -0.029 | 0.010 | 0.222 | downstream gene variant |
| rs55727654 | -0.0424 | 0.0345 | NA | NA | -0.027 | 0.014 | 0.151 | intron variant |
| rs5908 | 0.1216 | 0.0910 | 0.035 | 0.014 | -0.014 | 0.055 | 0.016 | missense variant |
| rs72633963 | 0.0982 | 0.0381 | NA | NA | -0.026 | 0.014 | 0.125 | non coding transcript exon variant |
| rs76475757 | -0.0167 | 0.0503 | -0.056 | 0.010 | -0.049 | 0.024 | 0.061 | intron variant |

**Appendix Table 2:** Genetic associations with pQTL, LDL-C and CHD in a $\pm 2.5$KB region around the *PCSK9* locus (clumped on $R^2 < 0.80$). GWAS data was extracted from Suhre[5], GLGC[2], and CardiogramPlusC4D[3, 4].

| rsID | Mean difference pQTL | SE pQTL | Mean difference per SD LDL | SE per SD LDL | logOR CHD | SE CHD | Allele frequency | Worst consequence |
|---|---|---|---|---|---|---|---|---|
| rs10465832 | NA | NA | -0.037 | 0.010 | -0.018 | 0.019 | 0.061 | intron variant |
| rs10888896 | NA | NA | -0.043 | 0.005 | -0.008 | 0.013 | 0.719 | intron variant |
| rs11206514 | NA | NA | -0.051 | 0.004 | -0.013 | 0.011 | 0.602 | intron variant |

**Appendix Table 2:** Genetic associations with pQTL, LDL-C and CHD in a $\pm 2.5$KB region around the *PCSK9* locus (clumped on $R^2 < 0.80$). GWAS data was extracted from Suhre[5], GLGC[2], and CardiogramPlusC4D[3, 4]. *(continued)*

| rsID | Mean difference pQTL | SE pQTL | Mean difference per SD LDL | SE per SD LDL | logOR CHD | SE CHD | Allele frequency | Worst consequence |
|---|---|---|---|---|---|---|---|---|
| rs11206517 | NA | NA | -0.063 | 0.014 | -0.054 | 0.027 | 0.035 | intron variant |
| rs11583680 | NA | NA | 0.034 | 0.006 | 0.037 | 0.017 | 0.134 | missense variant |
| rs11591147 | -0.642 | 0.153 | 0.497 | 0.018 | 0.355 | 0.069 | 0.021 | missense variant |
| rs11800243 | NA | NA | 0.014 | 0.014 | 0.044 | 0.030 | 0.053 | intron variant |
| rs12067569 | NA | NA | -0.088 | 0.010 | -0.038 | 0.023 | 0.030 | intron variant |
| rs13312 | NA | NA | -0.030 | 0.007 | 0.004 | 0.013 | 0.806 | 3 prime UTR variant |
| rs17111503 | NA | NA | -0.066 | 0.004 | -0.040 | 0.011 | 0.243 | upstream gene variant |
| rs2479409 | NA | NA | 0.064 | 0.004 | 0.037 | 0.010 | 0.680 | upstream gene variant |
| rs2483205 | NA | NA | 0.051 | 0.005 | 0.022 | 0.012 | 0.450 | splice region variant |
| rs2495477 | NA | NA | 0.064 | 0.005 | 0.036 | 0.012 | 0.400 | splice region variant |
| rs2495481 | NA | NA | 0.008 | 0.009 | 0.028 | 0.022 | 0.949 | intron variant |
| rs28385708 | NA | NA | 0.009 | 0.014 | 0.024 | 0.036 | 0.050 | intron variant |
| rs41294821 | NA | NA | 0.030 | 0.020 | 0.034 | 0.036 | 0.026 | intron variant |
| rs4927193 | NA | NA | 0.035 | 0.006 | 0.040 | 0.016 | 0.133 | intron variant |
| rs499718 | NA | NA | -0.036 | 0.008 | -0.025 | 0.014 | 0.824 | intron variant |
| rs505151 | 0.138 | 0.114 | NA | NA | 0.036 | 0.027 | NA | missense variant |
| rs529787 | NA | NA | 0.055 | 0.005 | 0.001 | 0.015 | 0.224 | intron variant |
| rs557435 | NA | NA | -0.062 | 0.007 | -0.001 | 0.014 | 0.799 | intron variant |
| rs572512 | NA | NA | -0.048 | 0.005 | -0.026 | 0.013 | 0.354 | non coding transcript exon variant |

**Appendix Table 2:** Genetic associations with pQTL, LDL-C and CHD in a $\pm 2.5$KB region around the *PCSK9* locus (clumped on $R^2 < 0.80$). GWAS data was extracted from Suhre[5], GLGC[2], and CardiogramPlusC4D[3, 4]. *(continued)*

| rsID | Mean difference pQTL | SE pQTL | Mean difference per SD LDL | SE per SD LDL | logOR CHD | SE CHD | Allele frequency | Worst consequence |
|---|---|---|---|---|---|---|---|---|
| rs585131 | NA | NA | -0.064 | 0.005 | -0.015 | 0.012 | 0.821 | intron variant |
| rs625619 | NA | NA | -0.042 | 0.005 | -0.008 | 0.012 | 0.556 | intron variant |
| rs630431 | NA | NA | -0.035 | 0.004 | -0.012 | 0.010 | 0.707 | intron variant |
| rs644000 | NA | NA | 0.060 | 0.006 | 0.022 | 0.012 | 0.362 | intron variant |
| rs662145 | NA | NA | -0.005 | 0.005 | 0.003 | 0.011 | 0.767 | 3 prime UTR variant |
| rs74700387 | NA | NA | 0.033 | 0.020 | 0.023 | 0.031 | 0.021 | intron variant |
| rs7552841 | NA | NA | -0.037 | 0.004 | -0.019 | 0.011 | 0.369 | intron variant |

**Appendix Table 3:** Genetic associations with LDL-C and CHD in a $\pm 2.5$KB region around the *NPC1L1* locus (clumped on $R^2 < 0.80$). GWAS data was extracted from GLGC[2], and CardiogramPlusC4D[3, 4].

| rsID | Mean difference per SD LDL | SE per SD LDL | logOR CHD | SE CHD | Allele frequency | Worst consequence |
|---|---|---|---|---|---|---|
| rs10264715 | -0.022 | 0.004 | -0.007 | 0.012 | 0.209 | synonymous variant |
| rs11763759 | 0.038 | 0.007 | 0.024 | 0.013 | 0.289 | intron variant |
| rs17655652 | 0.028 | 0.004 | 0.029 | 0.012 | 0.294 | upstream gene variant |
| rs2072183 | -0.039 | 0.005 | -0.004 | 0.013 | 0.234 | synonymous variant |
| rs2073547 | -0.048 | 0.005 | -0.004 | 0.014 | 0.195 | upstream gene variant |
| rs2073548 | -0.014 | 0.012 | 0.006 | 0.023 | 0.054 | upstream gene variant |
| rs217406 | -0.039 | 0.005 | -0.015 | 0.016 | 0.173 | intron variant |
| rs217420 | -0.022 | 0.004 | -0.007 | 0.013 | 0.227 | intron variant |

**Appendix Table 3:** Genetic associations with LDL-C and CHD in a $\pm 2.5$KB region around the *NPC1L1* locus (clumped on $R^2 < 0.80$). GWAS data was extracted from GLGC[2], and CardiogramPlusC4D[3, 4]. *(continued)*

| rsID | Mean difference per SD LDL | SE per SD LDL | logOR CHD | SE CHD | Allele frequency | Worst consequence |
|---|---|---|---|---|---|---|
| rs217426 | 0.005 | 0.011 | -0.028 | 0.027 | 0.029 | intron variant |
| rs217433 | -0.020 | 0.005 | -0.008 | 0.011 | 0.179 | intron variant |
| rs217437 | -0.008 | 0.004 | -0.003 | 0.012 | 0.626 | downstream gene variant |
| rs35349497 | 0.018 | 0.009 | 0.008 | 0.022 | 0.091 | intron variant |
| rs41279627 | 0.026 | 0.013 | 0.048 | 0.034 | 0.047 | intron variant |
| rs41279633 | -0.052 | 0.007 | -0.008 | 0.016 | 0.143 | 5 prime UTR variant |
| rs4720470 | 0.002 | 0.010 | 0.003 | 0.022 | 0.067 | intron variant |
| rs73107472 | -0.009 | 0.022 | -0.031 | 0.049 | 0.014 | intron variant |

**Appendix Table 4:** Genetic associations with pQTL, HDL-C and CHD in a $\pm 2.5$KB region around the *CETP* locus (clumped on $R^2 < 0.80$). GWAS data was extracted from Blauw [6], GLGC[2], and CardiogramPlusC4D[3, 4].

| rsID | Mean difference pQTL | SE pQTL | Mean difference per SD HDL | SE per SD HDL | logOR CHD | SE CHD | Allele frequency | Worst consequence |
|---|---|---|---|---|---|---|---|---|
| rs11076174 | -0.236 | 0.029 | 0.180 | 0.008 | -0.056 | 0.020 | 0.108 | intron variant |
| rs11076175 | NA | NA | 0.254 | 0.005 | -0.034 | 0.014 | 0.206 | intron variant |
| rs117040820 | 0.356 | 0.062 | -0.181 | 0.022 | 0.006 | 0.061 | 0.013 | intron variant |
| rs118146573 | -0.379 | 0.023 | 0.262 | 0.007 | -0.023 | 0.017 | 0.122 | intron variant |
| rs12597002 | NA | NA | 0.085 | 0.004 | -0.017 | 0.012 | 0.280 | intron variant |
| rs12708974 | NA | NA | -0.008 | 0.006 | 0.011 | 0.016 | 0.134 | intron variant |
| rs12708980 | NA | NA | 0.016 | 0.004 | -0.002 | 0.009 | 0.351 | intron variant |
| rs12720873 | NA | NA | -0.095 | 0.014 | -0.019 | 0.038 | 0.026 | intron variant |
| rs12720898 | NA | NA | -0.022 | 0.013 | 0.017 | 0.021 | 0.066 | intron variant |

**Appendix Table 4:** Genetic associations with pQTL, HDL-C and CHD in a $\pm 2.5$KB region around the *CETP* locus (clumped on $R^2 < 0.80$). GWAS data was extracted from Blauw [6], GLGC[2], and CardiogramPlusC4D[3, 4]. *(continued)*

| rsID | Mean difference pQTL | SE pQTL | Mean difference per SD HDL | SE per SD HDL | logOR CHD | SE CHD | Allele frequency | Worst consequence |
|---|---|---|---|---|---|---|---|---|
| rs12720917 | 0.140 | 0.024 | -0.098 | 0.006 | 0.012 | 0.018 | 0.140 | upstream gene variant |
| rs1532624 | NA | NA | -0.204 | 0.004 | 0.027 | 0.011 | 0.426 | intron variant |
| rs17231506 | NA | NA | -0.238 | 0.004 | 0.027 | 0.011 | 0.291 | upstream gene variant |
| rs17231534 | NA | NA | 0.016 | 0.012 | 0.015 | 0.024 | 0.045 | 5 prime UTR variant |
| rs17245715 | NA | NA | -0.001 | 0.008 | 0.001 | 0.017 | 0.124 | upstream gene variant |
| rs1800775 | 0.267 | 0.014 | -0.202 | 0.004 | 0.031 | 0.010 | 0.488 | upstream gene variant |
| rs1800776 | NA | NA | 0.057 | 0.010 | -0.007 | 0.024 | 0.065 | upstream gene variant |
| rs1801706 | NA | NA | -0.064 | 0.008 | -0.002 | 0.014 | 0.193 | 3 prime UTR variant |
| rs1864163 | -0.321 | 0.017 | 0.225 | 0.004 | -0.030 | 0.012 | 0.270 | intron variant |
| rs289714 | NA | NA | -0.214 | 0.005 | 0.014 | 0.014 | 0.798 | intron variant |
| rs289715 | NA | NA | 0.134 | 0.006 | -0.060 | 0.050 | 0.878 | intron variant |
| rs289717 | NA | NA | 0.085 | 0.005 | -0.013 | 0.012 | 0.326 | intron variant |
| rs289719 | NA | NA | 0.113 | 0.004 | -0.023 | 0.011 | 0.670 | intron variant |
| rs289745 | NA | NA | -0.028 | 0.004 | -0.004 | 0.013 | 0.604 | upstream gene variant |
| rs4783961 | 0.133 | 0.015 | -0.100 | 0.004 | 0.022 | 0.011 | 0.469 | upstream gene variant |
| rs4783962 | NA | NA | -0.075 | 0.004 | 0.010 | 0.011 | 0.770 | upstream gene variant |
| rs5030708 | NA | NA | 0.079 | 0.016 | -0.014 | 0.037 | 0.034 | intron variant |
| rs5880 | NA | NA | 0.307 | 0.009 | -0.025 | 0.024 | 0.052 | missense variant |

**Appendix Table 4:** Genetic associations with pQTL, HDL-C and CHD in a $\pm 2.5$KB region around the *CETP* locus (clumped on $R^2 < 0.80$). GWAS data was extracted from Blauw [6], GLGC[2], and CardiogramPlusC4D[3, 4]. *(continued)*

| rsID | Mean difference pQTL | SE pQTL | Mean difference per SD HDL | SE per SD HDL | logOR CHD | SE CHD | Allele frequency | Worst consequence |
|---|---|---|---|---|---|---|---|---|
| rs5883 | NA | NA | -0.115 | 0.008 | -0.024 | 0.026 | 0.056 | synonymous variant |
| rs820299 | NA | NA | -0.064 | 0.004 | 0.005 | 0.011 | 0.637 | intron variant |
| rs9923854 | NA | NA | -0.084 | 0.010 | -0.022 | 0.019 | 0.127 | intron variant |
| rs9929488 | NA | NA | 0.176 | 0.004 | -0.024 | 0.012 | 0.289 | intron variant |
| rs9930761 | NA | NA | -0.062 | 0.008 | -0.014 | 0.023 | 0.073 | intron variant |

**Appendix Table 5:** Number of variants with lipids or pQTL, and CHD data in a $\pm 2.5$KB region around each locus; stratified by type of mutation (after clumping on $R^2 < 0.80$)

| | ***HMGCR*** | ***PCSK9*** | ***NPC1L1*** | ***CETP*** |
|---|---|---|---|---|
| Total number of variants | 26 | 29 | 16 | 32 |
| Number of pQTL variants | 21 | 2 | 0 | 7 |
| Number of GLGC variants | 14 | 28 | 16 | 32 |
| 3 prime UTR variant | 2 | 2 | 0 | 1 |
| 5 prime UTR variant | 1 | 0 | 1 | 1 |
| Downstream gene variant | 1 | 0 | 1 | 0 |
| Intron variant | 17 | 19 | 9 | 20 |
| Missense variant | 1 | 3 | 0 | 1 |
| Non coding transcript exon variant | 3 | 1 | 0 | 0 |
| Upstream gene variant | 1 | 2 | 3 | 8 |
| Splice region variant | 0 | 2 | 0 | 0 |
| Synonymous variant | 0 | 0 | 2 | 1 |

**Appendix Table 6:** The *HMGCR* lipids ~ CHD effect, comparing the GLS estimates to functional specific MR estimates

| *Type of variant selection* | *nSNPs* | *Fixed effect* | *Random effects* |
|---|---|---|---|
| GLS | 10 | 1.68 (1.35;2.10) | 1.68 (1.35;2.10) |
| 3 prime UTR variant | 2 | 1.68 (1.32;2.12) | 1.68 (1.20;2.34) |
| intron variant | 5 | 1.80 (1.30;2.48) | 1.80 (1.30;2.48) |
| non coding transcript exon variant | 1 | 2.99 (0.13;71.42) | NA |
| downstream gene variant | 1 | 1.61 (1.17;2.22) | NA |
| missense variant | 1 | 0.67 (0.03;14.02) | NA |
| upstream gene variant | NA | NA | NA |
| splice region variant | NA | NA | NA |
| 5 prime UTR variant | NA | NA | NA |
| synonymous variant | NA | NA | NA |
| Miss & splice | 1 | 0.67 (0.03;14.02) | NA |
| 3- & 5-UTR & intron & nc exon | 8 | 1.66 (1.32;2.08) | 1.66 (1.32;2.08) |
| Down & upstream | 1 | 1.61 (1.17;2.22) | NA |

**Appendix Table 7:** The *PCSK9* lipids ~ CHD effect, comparing the GLS estimates to functional specific MR estimates

| *Type of variant selection* | *nSNPs* | *Fixed effect* | *Random effects* |
|---|---|---|---|
| GLS | 21 | 1.83 (1.58;2.12) | 1.83 (1.55;2.17) |
| 3 prime UTR variant | NA | NA | NA |
| intron variant | 16 | 1.53 (1.25;1.88) | 1.53 (1.24;1.90) |
| non coding transcript exon variant | 1 | 1.71 (1.02;2.88) | NA |
| downstream gene variant | NA | NA | NA |
| missense variant | 1 | 2.04 (1.56;2.67) | NA |
| upstream gene variant | 1 | 1.78 (1.31;2.41) | NA |
| splice region variant | 2 | 1.70 (1.21;2.40) | 1.70 (1.21;2.40) |
| 5 prime UTR variant | NA | NA | NA |
| synonymous variant | NA | NA | NA |
| Miss & splice | 3 | 1.91 (1.54;2.36) | 1.91 (1.54;2.36) |
| 3- & 5-UTR & intron & nc exon | 17 | 1.64 (1.35;2.00) | 1.64 (1.31;2.06) |
| Down & upstream | 1 | 1.78 (1.31;2.41) | NA |

**Appendix Table 8:** The *NPC1L1* lipids ~ CHD effect, comparing the GLS estimates to functional specific MR estimates

| *Type of variant selection* | *nSNPs* | *Fixed effect* | *Random effects* |
|---|---|---|---|
| GLS | 11 | 1.43 (0.89;2.29) | 1.43 (0.89;2.29) |
| 3 prime UTR variant | NA | NA | NA |
| intron variant | 6 | 1.67 (0.80;3.50) | 1.67 (0.80;3.50) |
| non coding transcript exon variant | NA | NA | NA |
| downstream gene variant | 1 | 1.43 (0.07;28.14) | NA |
| missense variant | NA | NA | NA |
| upstream gene variant | 3 | 1.35 (0.82;2.22) | 1.35 (0.63;2.86) |
| splice region variant | NA | NA | NA |
| 5 prime UTR variant | NA | NA | NA |
| synonymous variant | 1 | 1.38 (0.48;3.91) | NA |
| Miss & splice | NA | NA | NA |
| 3- & 5-UTR & intron & nc exon | 6 | 1.67 (0.80;3.50) | 1.67 (0.80;3.50) |
| Down & upstream | 4 | 1.35 (0.82;2.22) | 1.35 (0.72;2.52) |

**Appendix Table 9:** The *CETP* lipids ~ CHD effect, comparing the GLS estimates to functional specific MR estimates

| *Type of variant selection* | *nSNPs* | *Fixed effect* | *Random effects* |
|---|---|---|---|
| GLS | 26 | 0.84 (0.80;0.90) | 0.84 (0.78;0.91) |
| 3 prime UTR variant | 1 | 1.04 (0.67;1.61) | NA |
| intron variant | 17 | 0.83 (0.77;0.88) | 0.83 (0.75;0.90) |
| non coding transcript exon variant | NA | NA | NA |
| downstream gene variant | NA | NA | NA |
| missense variant | 1 | 0.92 (0.79;1.08) | NA |
| upstream gene variant | 5 | 0.92 (0.84;1.00) | 0.92 (0.84;1.00) |
| splice region variant | NA | NA | NA |
| 5 prime UTR variant | 1 | 2.56 (0.14;47.57) | NA |
| synonymous variant | 1 | 1.24 (0.79;1.94) | NA |
| Miss & splice | 1 | 0.92 (0.79;1.08) | NA |
| 3- & 5-UTR & intron & nc exon | 19 | 0.84 (0.79;0.89) | 0.84 (0.76;0.92) |

| | | | |
|---|---|---|---|
| Down & upstream | 5 | 0.92 (0.84;1.00) | 0.92 (0.84;1.00) |

**Appendix Table 10:** Number of variants with eQTL and CHD data in a $\pm 2.5$KB region around each locus; stratified by tissue (after clumping on $R^2 < 0.60$). GWAS data was extracted from GTEx [7].

| | ***HMGCR*** | ***PCSK9*** | ***NPC1L1*** | ***CETP*** |
|---|---|---|---|---|
| Median (Q1;Q3) number of variants across tissues | 1 (0; 28.25) | 4 (1; 6) | 2 (1; 19.5) | 4.5 (2.75; 7.25) |
| Adipose Subcutaneous | 33 | 1 | 1 | 8 |
| Adipose Visceral Omentum | 1 | 8 | 2 | 7 |
| Adrenal Gland | 21 | 2 | 18 | 4 |
| Artery Aorta | 1 | 70 | 2 | 3 |
| Artery Coronary | 20 | 1 | 1 | 4 |
| Artery Tibial | 0 | 3 | 45 | 53 |
| Brain Amygdala | 33 | 0 | 0 | 58 |
| Brain Anterior cingulate cortex BA24 | 1 | 1 | 0 | 1 |
| Brain Caudate basal ganglia | 45 | 0 | 2 | 2 |
| Brain Cerebellar Hemisphere | 0 | 5 | 1 | 1 |
| Brain Cerebellum | 46 | 11 | 37 | 54 |
| Brain Cortex | 1 | 7 | 31 | 2 |
| Brain Frontal Cortex BA9 | 61 | 2 | 2 | 2 |
| Brain Hippocampus | 1 | 0 | 0 | 1 |
| Brain Hypothalamus | 1 | 3 | 0 | 45 |
| Brain Nucleus accumbens basal ganglia | 1 | 4 | 0 | 53 |
| Brain Putamen basal ganglia | 1 | 0 | 1 | 40 |
| Brain Spinal cord cervical c-1 | 32 | 3 | 1 | 3 |

**Appendix Table 10:** Number of variants with eQTL and CHD data in a $\pm 2.5$KB region around each locus; stratified by tissue (after clumping on $R^2 < 0.60$). GWAS data was extracted from GTEx [7]. *(continued)*

| | ***HMGCR*** | ***PCSK9*** | ***NPC1L1*** | ***CETP*** |
|---|---|---|---|---|
| Brain Substantia nigra | 40 | 48 | 24 | 4 |
| Breast Mammary Tissue | 26 | 2 | 3 | 3 |
| Cells EBV-transformed lymphocytes | 48 | 0 | 47 | 9 |
| Cells Transformed fibroblasts | 0 | 5 | 2 | 9 |
| Colon Sigmoid | 45 | 4 | 1 | 1 |
| Colon Transverse | 1 | 4 | 0 | 7 |
| Esophagus Gastroesophageal Junction | 40 | 4 | 27 | 6 |
| Esophagus Mucosa | 1 | 1 | 1 | 3 |
| Esophagus Muscularis | 0 | 1 | 1 | 1 |
| Heart Atrial Appendage | 0 | 0 | 1 | 6 |
| Heart Left Ventricle | 0 | 0 | 1 | 1 |
| Liver | 36 | 4 | 41 | 7 |
| Lung | 1 | 6 | 1 | 7 |
| Minor Salivary Gland | 0 | 39 | 2 | 4 |
| Muscle Skeletal | 1 | 0 | 2 | 7 |
| Nerve Tibial | 0 | 6 | 2 | 3 |
| Ovary | 0 | 62 | 36 | 2 |
| Pancreas | 1 | 5 | 2 | 4 |
| Pituitary | 1 | 47 | 39 | 6 |
| Prostate | 0 | 1 | 1 | 9 |
| Skin Not Sun Exposed Suprapubic | 0 | 5 | 1 | 1 |
| Skin Sun Exposed Lower leg | 1 | 6 | 46 | 3 |

**Appendix Table 10:** Number of variants with eQTL and CHD data in a $\pm 2.5$KB region around each locus; stratified by tissue (after clumping on $R^2 < 0.60$). GWAS data was extracted from GTEx [7]. *(continued)*

| | ***HMGCR*** | ***PCSK9*** | ***NPC1L1*** | ***CETP*** |
|---|---|---|---|---|
| Small Intestine Terminal Ileum | 0 | 4 | 53 | 8 |
| Spleen | 27 | 5 | 0 | 3 |
| Stomach | 1 | 3 | 47 | 7 |
| Testis | 1 | 12 | 2 | 5 |
| Thyroid | 1 | 0 | 1 | 5 |
| Uterus | 38 | 4 | 1 | 2 |
| Vagina | 1 | 4 | 2 | 64 |
| Whole Blood | 0 | 11 | 0 | 6 |

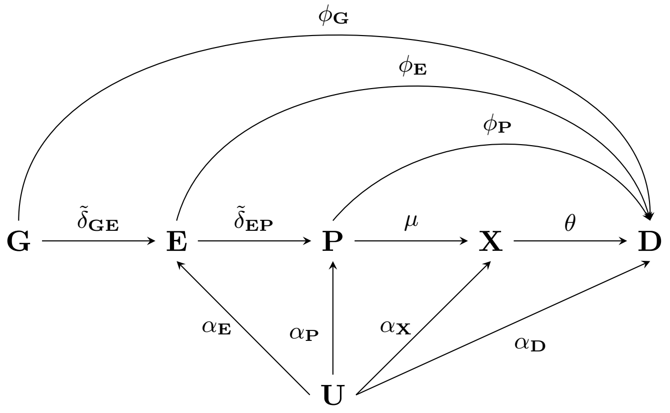


**Appendix Figure 1:** A directed acyclic graph depicting a Mendelian randomization study where eQTL ($\mathbf{G} \rightarrow \mathbf{E}$), downstream biomarker ($\mathbf{G} \rightarrow \mathbf{X}$), and pQTL ($\mathbf{G} \rightarrow \mathbf{P}$) weights would *all* results in a valid MR test of the causal $\mathbf{P} \rightarrow \mathbf{D}$ pathway; conditional on $\phi_{\mathbf{G}} = \phi_{\mathbf{E}} = 0$

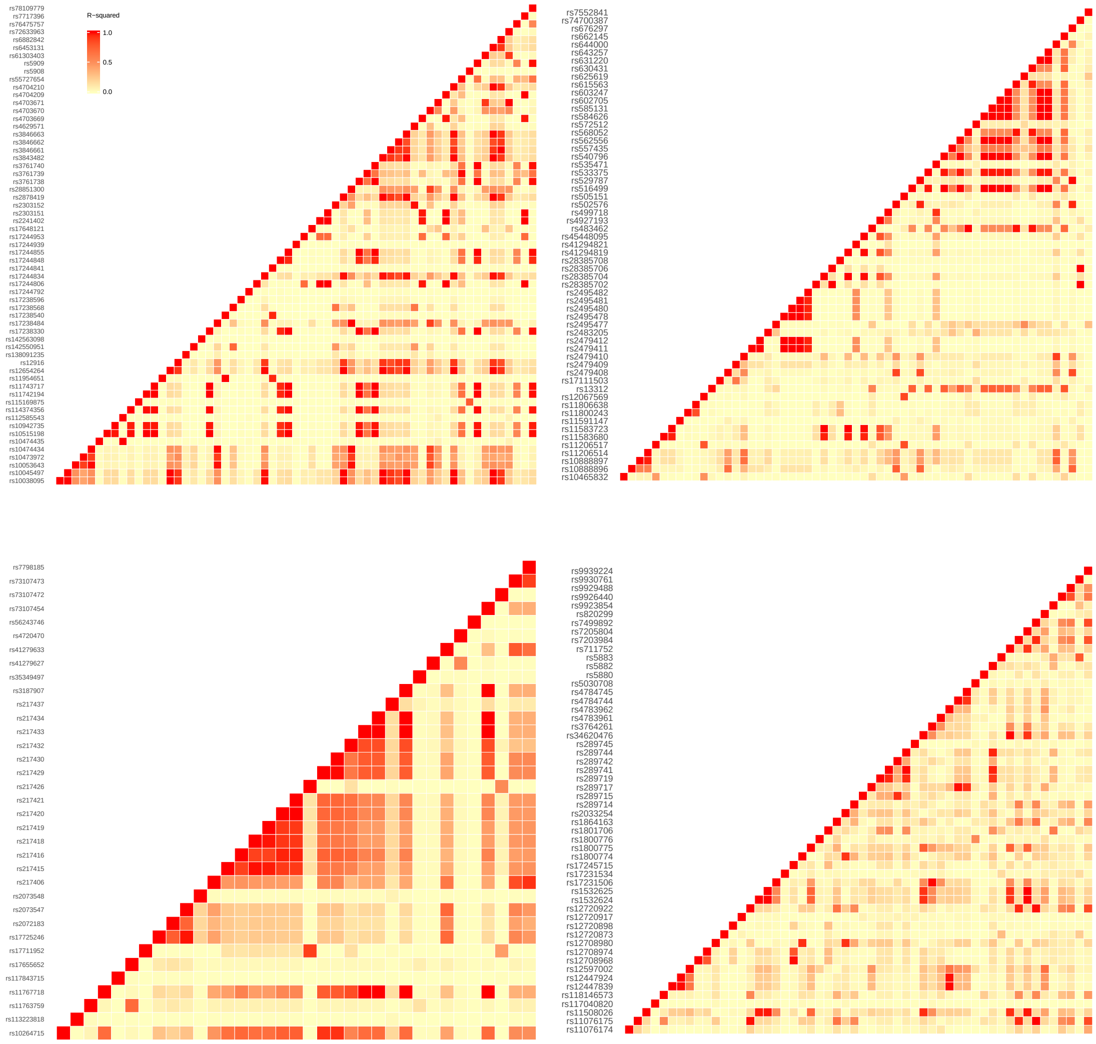


**Appendix Figure 2:** Linkage disequilibrium heatmaps of variants with aggregated data on lipids or pQTL associations. Pairwise LD was based on the "EUR" 1000 genomes panel [8]. Left top panel *HMGCR* locus, top right panel *PCSK9* locus, bottom left *NPC1L1* locus, and bottom right *CETP* locus; y-axis labels: rsIDs.

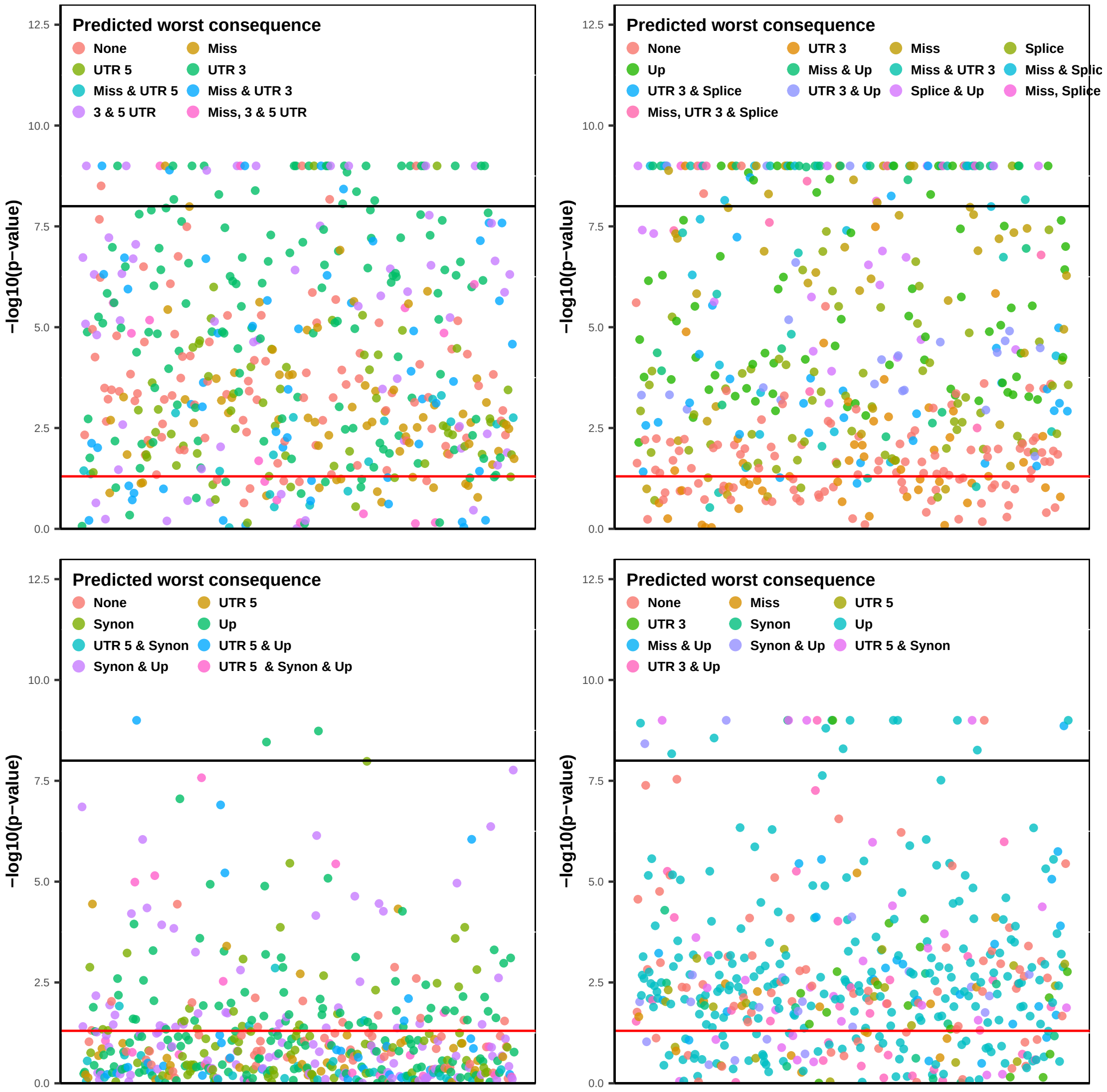


**Appendix Figure 3:** A Manhattan plot of 500 repeat-sampled MR analysis using a 4 instruments genetic score. Instruments were taken from the *HMGCR* locus (top left), the *PCSK9* locus (top right panel), *NPC1L1* locus (bottom left), and *CETP* locus (bottom right). Horizontal lines indicate p-value thresholds for 0.05 and $10^{-8}$, p-values were truncated at $10^{-9}$. Multiplicative random-effects standard error estimates were corrected for LD using the "EUR" 1000 genomes panel [8] and the estimator proposed in [9, 10].

#### Simulation study on the influence of null-variants

Instrument selection in Mendelian randomization (MR) revolves around selecting genetic variants that predict a risk factor, which in turn may affect disease. As with any selection, one will unavoidably make mistakes and exclude variants that are truly predictive of the intermediate risk factors, as well as include variants that do not predict the risk factor; *null-variants*.

To explore the influence of erroneous inclusion of null-variants we designed the following simulation study, where genetic and phenotypic information was simulated for 200 subjects. For each subject we simulated 100 genetic variants with a minor allele frequency of 0.30. The intermediate risk factor was generated based on an unobserved confounder $\mathbf{U} \sim N(0, 2)$ and the genetic variants $\mathbf{G}$:

$$x = u + \sum g\tilde{\delta\mu} + \varepsilon,$$

Where $\varepsilon \sim N(0, 1)$ and $\tilde{\delta\mu} \sim U(0.5, 4)$. Null-variants were generated by setting $\tilde{\delta\mu} = 0$ for $p$ many variants, where $P$ followed a Bernoulli distribution with probability $\{0.0, 0.1, \ldots, 0.9\}$. Subsequently the outcome phenotype was generated based on:

$$q = u + x\theta,$$
$$\mathbf{D} \sim Bernoulli(q),$$

here $\theta = 0$, so there is no causal effect of the risk factor on the outcome. To simulate a two-sample MR, the above described algorithm was run twice, were the first dataset was used to determine the variant to risk factor effect estimates, and the second, independent, dataset was used to estimate the variant to outcome effect estimates; the simulation was repeated 2,000 times.

For each iteration, we selected *all* variants and compared this to an *oracle* selection strategy, only including truly associated instruments. MR-estimates (without Egger correction) based on both selection strategies were evaluated on bias and root mean squared error (RMSE: the square root of the squared bias plus variance of the MR point estimate).

**Appendix Table 11:** Simulation study on the influence of null-variants on drug-target MR results; bias (RMSE)

| Proportion of Null-variants | All variants | Oracle |
|---|---|---|
| 0.0 | 0.00<br>(0.010) | 0.00<br>(0.010) |
| 0.1 | -0.00<br>(0.011) | -0.00<br>(0.011) |
| 0.2 | -0.00<br>(0.012) | -0.00<br>(0.012) |
| 0.3 | 0.00<br>(0.013) | 0.00<br>(0.014) |
| 0.4 | 0.00<br>(0.014) | 0.00<br>(0.015) |
| 0.5 | 0.00<br>(0.015) | 0.00<br>(0.017) |
| 0.6 | -0.00<br>(0.017) | -0.00<br>(0.019) |
| 0.7 | 0.00<br>(0.020) | 0.00<br>(0.023) |
| 0.8 | -0.00<br>(0.024) | -0.00<br>(0.029) |
| 0.9 | -0.00<br>(0.034) | -0.00<br>(0.043) |

The simulation results show that 1) inclusion of null-variants does not increase bias, and 2) typically variance (RMSE) is similar to only including the truly associated instrument (Oracle selection). Essentially, assuming one has included strong instruments including some additional null-variants does not detract from the results. This behaviour is obviously very much desired, especially given the impossibility of perfectly differentiating between null-variants and truly associated variants.

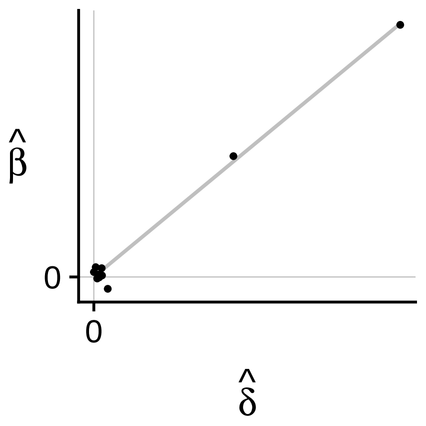


**Appendix Figure 4:** A hypothetical example showing null-variants do not influence the Mendelian randomization effect estimate (the slope). N.b. $\hat{\delta}$ values were generated based on $N\left(0, 0.01^2\right)$ and adding the constants $0.0, 0.2$, or $0.4$; $\hat{\beta} = 1.1\hat{\delta} + N\left(0, 0.01^2\right)$.

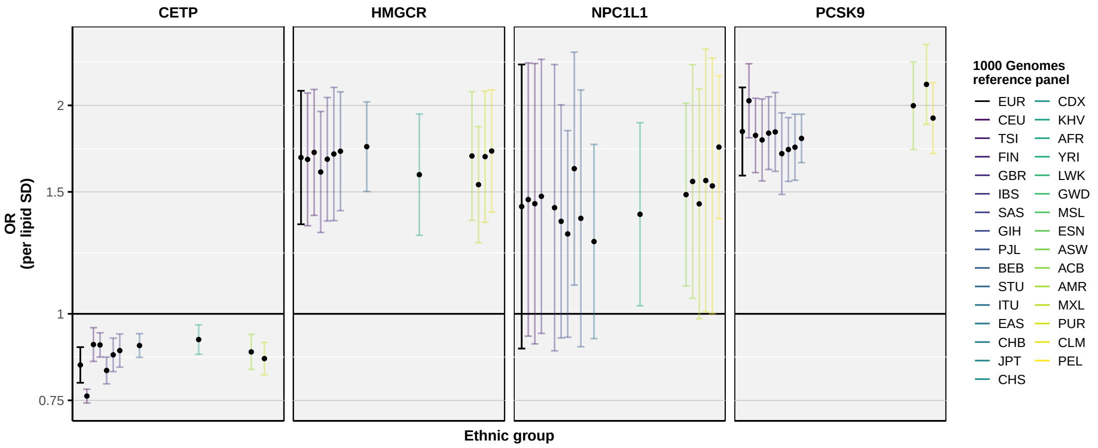


**Appendix Figure 5:** Mendelian randomization estimates of lipids weighted associations with CHD, applying LD corrections from different ethnic source populations. To highlight difference between 1000 genomes reference populations fixed effect standard errors were used. Similarly, the candidate variants were kept constant across the different 1000 genomes populations[8] by selecting variants with an R-squared clumping threshold of 0.60 as inferred from the EUR panel. Estimates are given as odd ratios (ORs) and 95% confidence intervals, respectively.

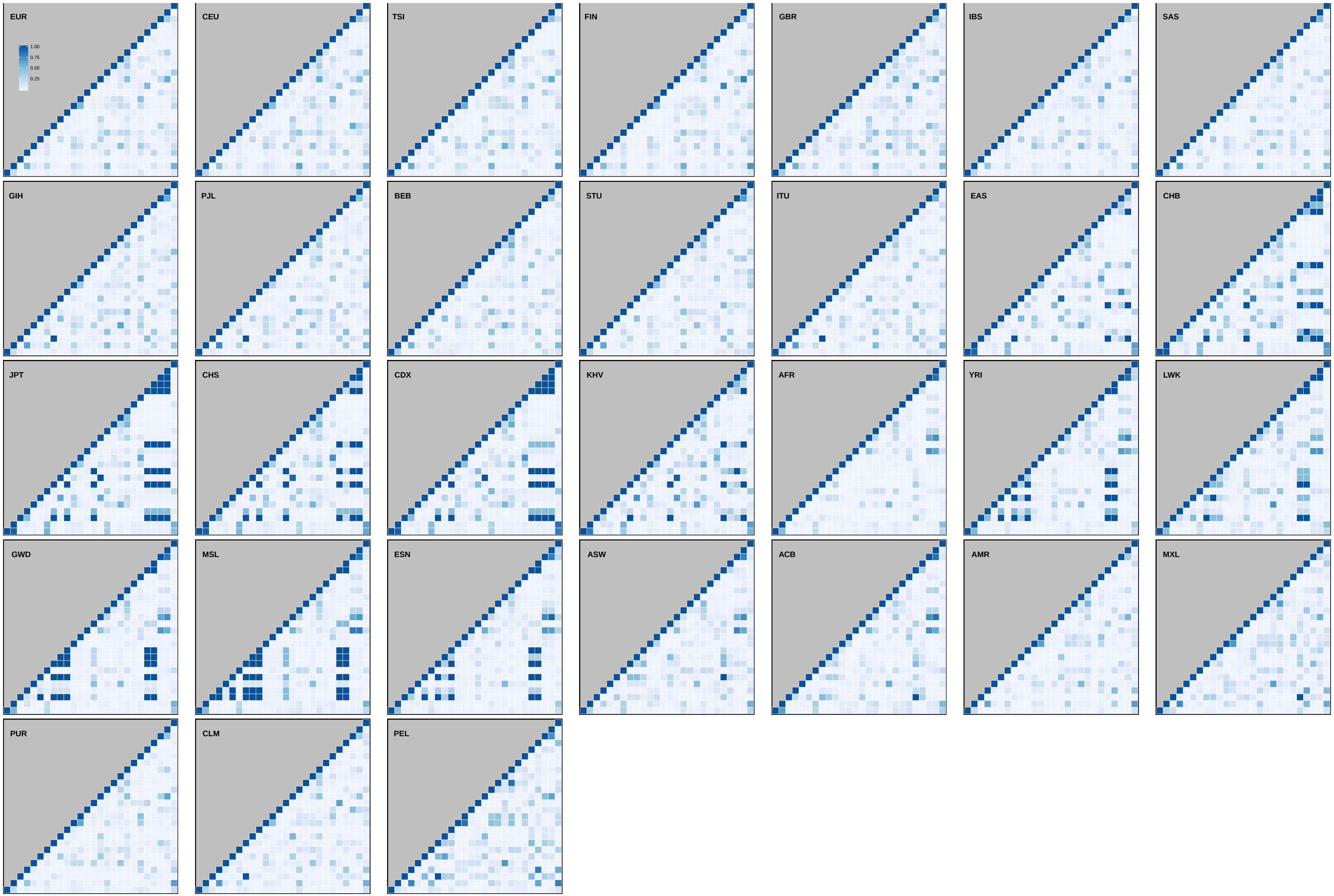


**Appendix Figure 6:** Linkage disequilibrium heatmaps from the *CETP* gene using the different 1000 genomes reference panel data[8]. Missing variants are depicted as row and column combinations with background colouring.

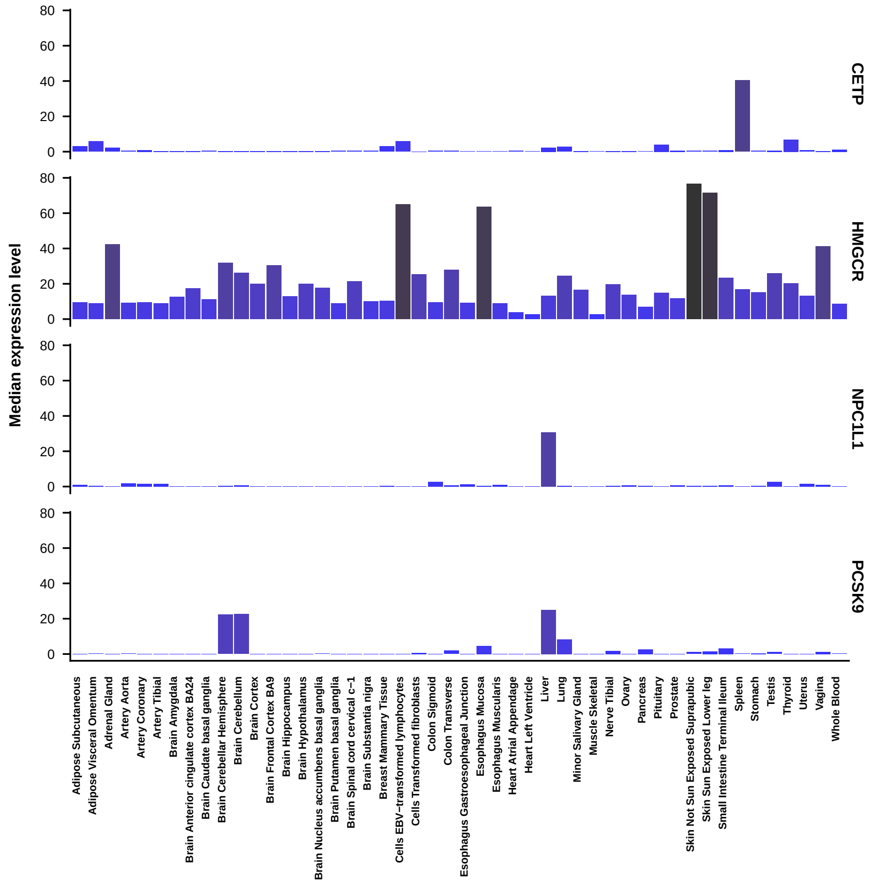


**Appendix Figure 7:** Tissue-specific expression level of 4 known drug-targets

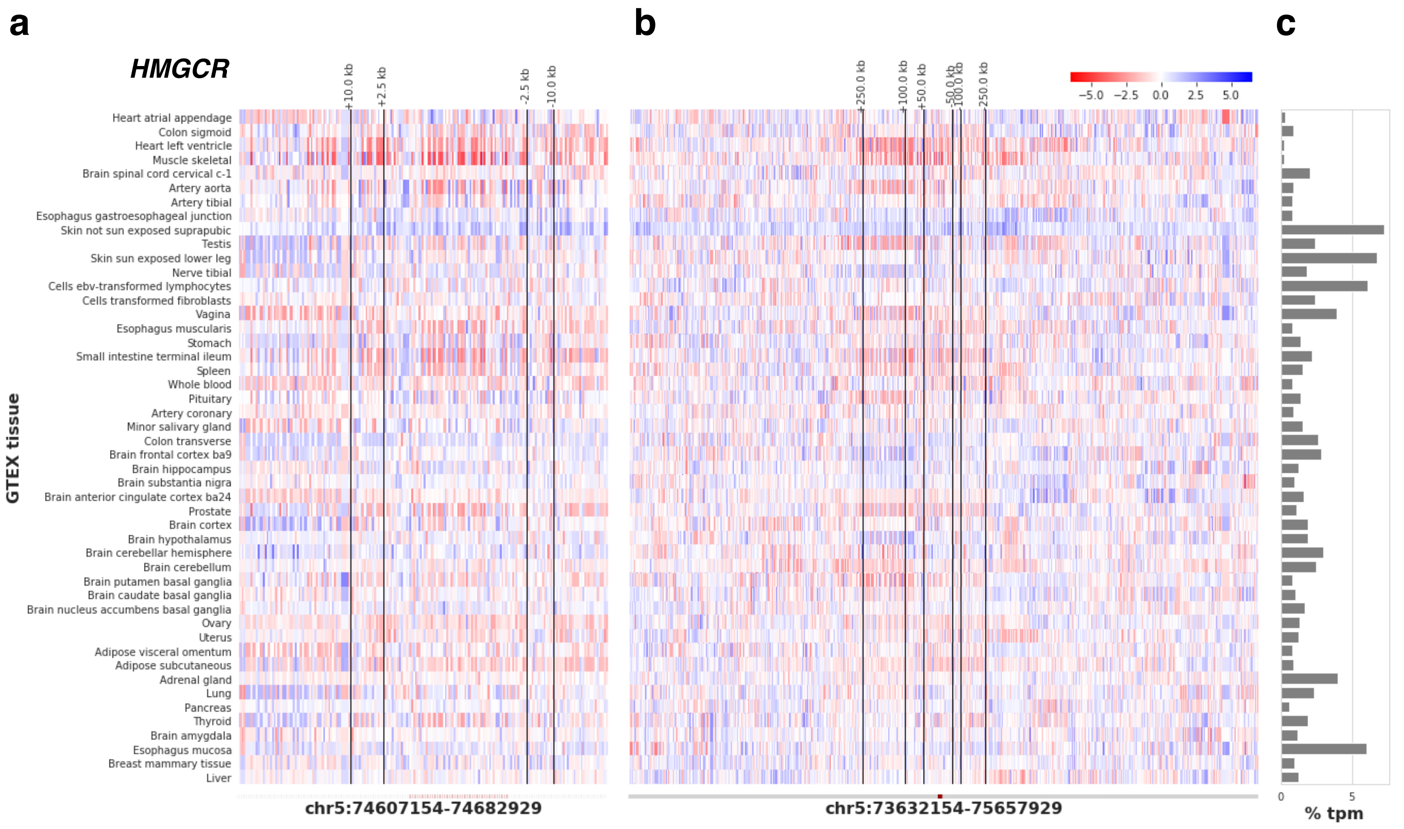


**Appendix Figure 8:** eQTL associations in *HMGCR* presented as z-statistics. x-axis indicates the genomic coordinates ±25 kbp (a) and 1Mbp (b), with the gene coloured in maroon. The relative gene expression in transcript per million (%tpm) is shown in c. The alleles are referenced to skin not sun exposed (tissue with the maximum % tpm).

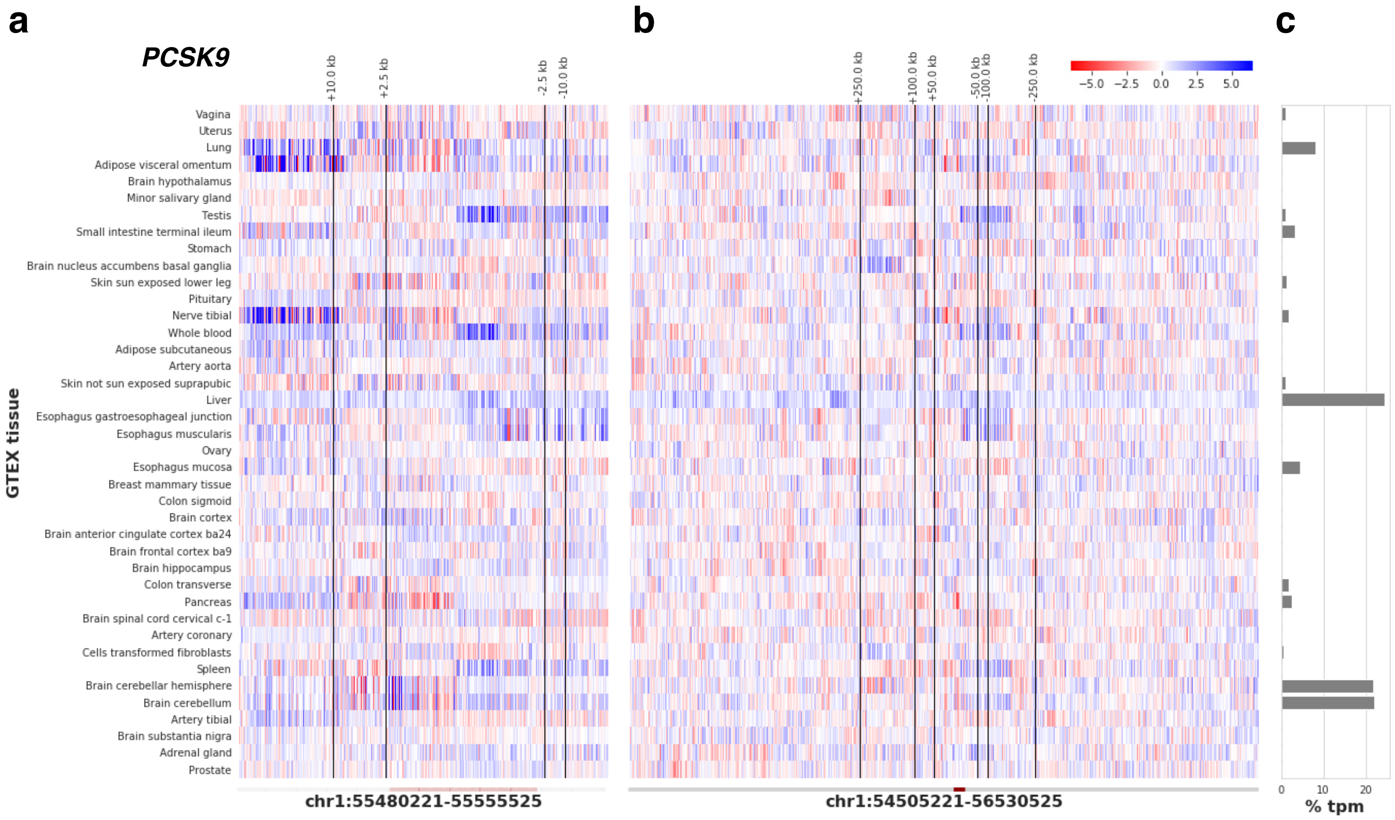


**Appendix Figure 9:** eQTL associations in *PCSK9* presented as z-statistics. x-axis indicates the genomic coordinates ±25 kbp (a) and 1Mbp (b), with the gene coloured in maroon. The relative gene expression in transcript per million (%tpm) is shown in c. The alleles are referenced to liver (tissue with the maximum % tpm).

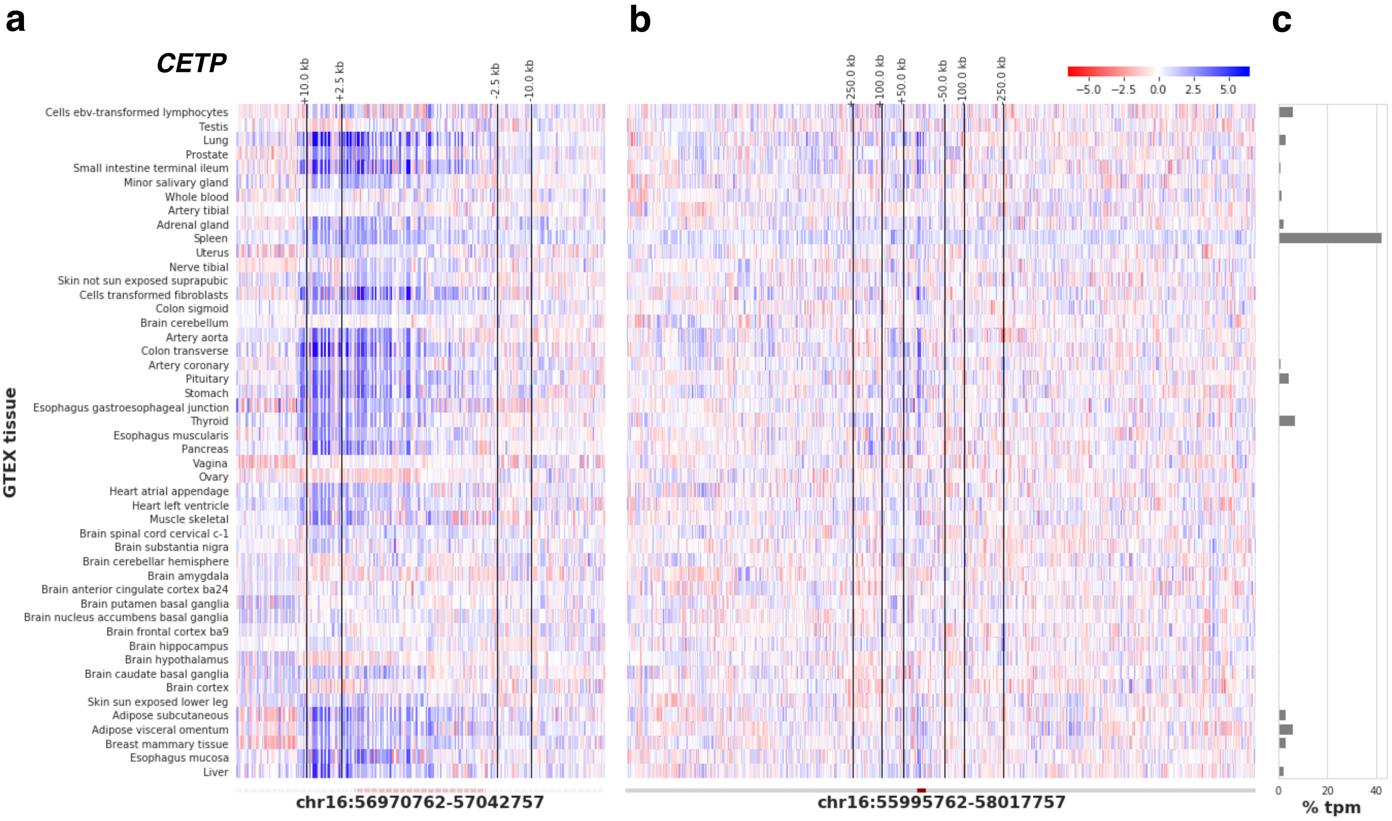


**Appendix Figure 10:** eQTL associations in *CETP* presented as z-statistics. x-axis indicates the genomic coordinates ±25 kbp (a) and 1Mbp (b), with the gene coloured in maroon. The relative gene expression in transcript per million (%tpm) is shown in c. The alleles are referenced to spleen (tissue with the maximum % tpm).

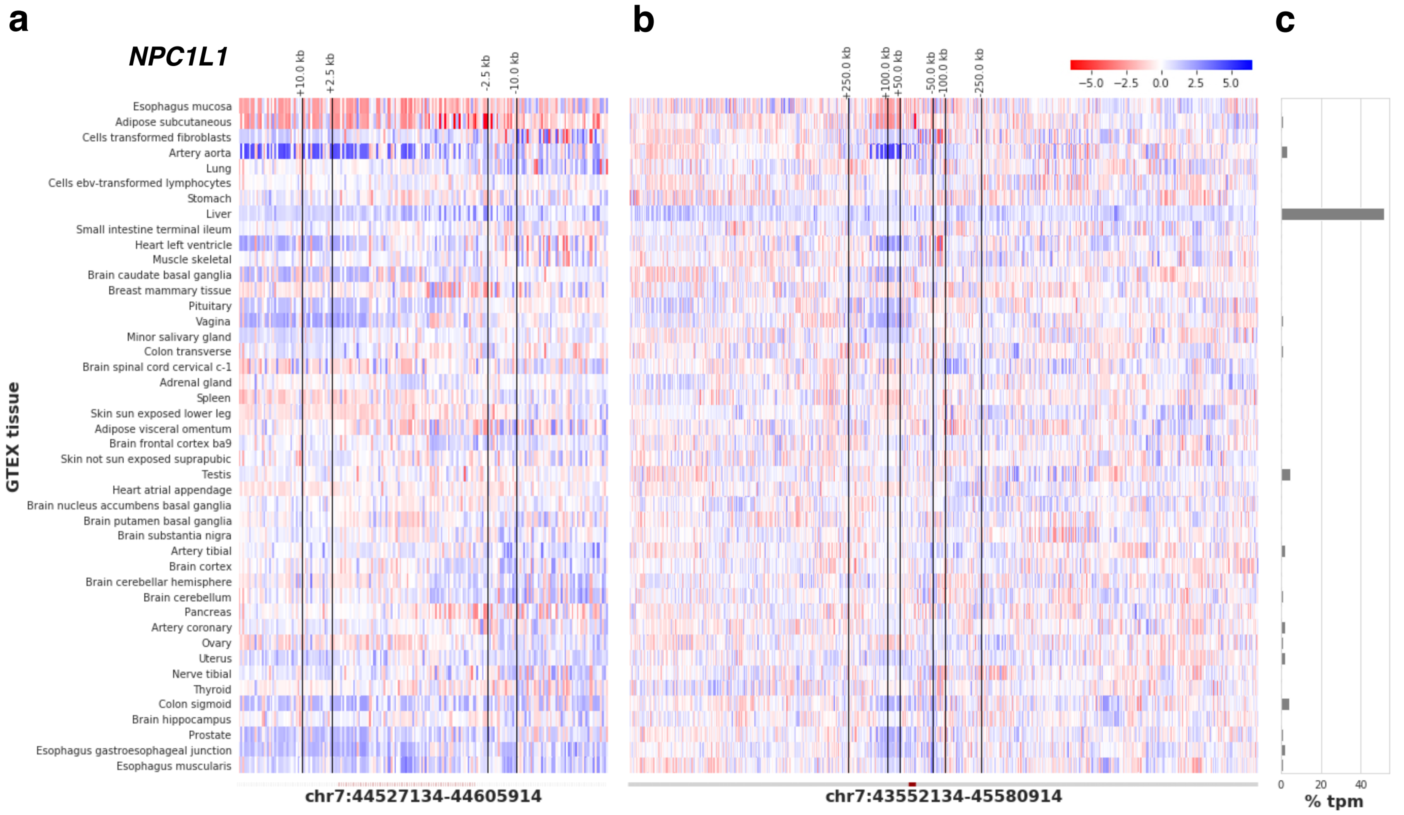


**Appendix Figure 11:** eQTL associations in *NPC1L1* presented as z-statistics. x-axis indicates the genomic coordinates ±25 kbp (a) and 1Mbp (b), with the gene coloured in maroon. The relative gene expression in transcript per million (%tpm) is shown in c. The alleles are referenced to liver (tissue with the maximum % tpm).

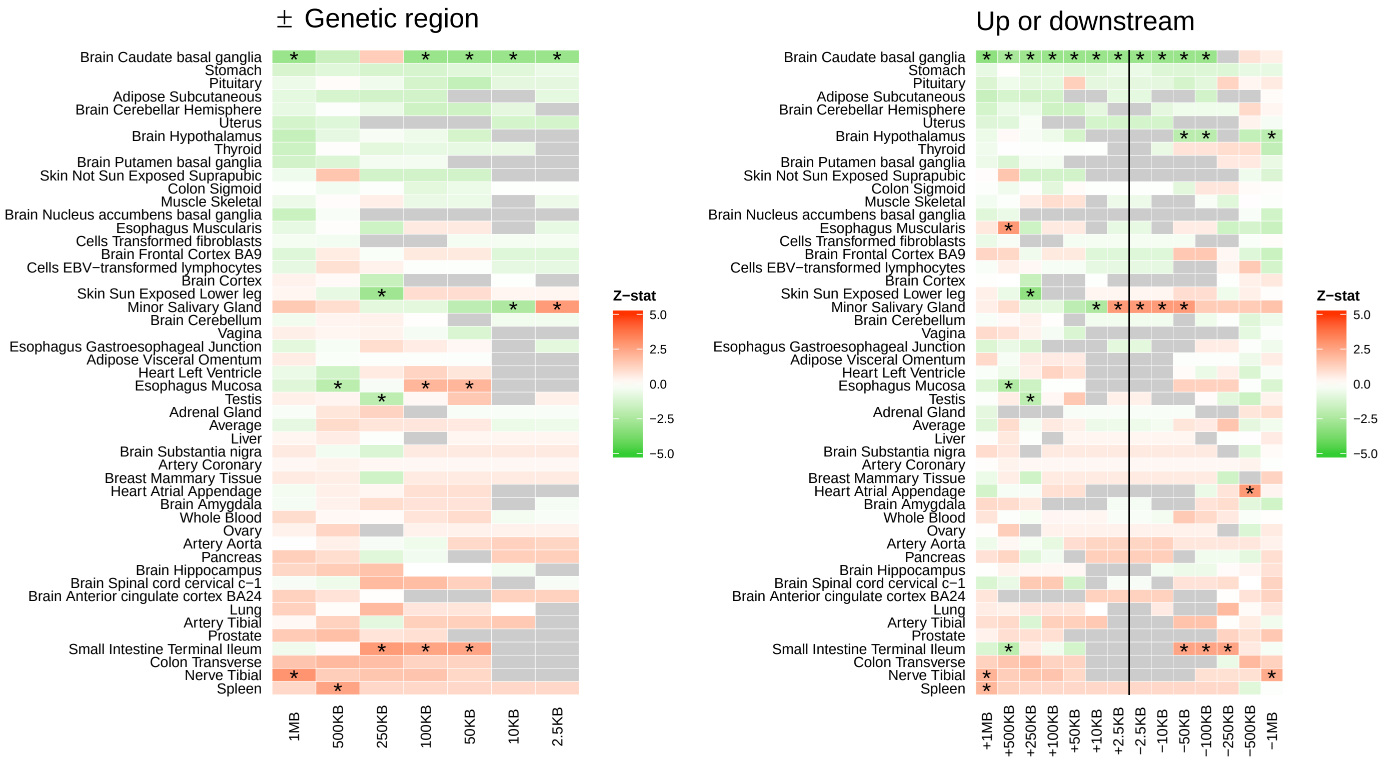


**Appendix Figure 12:** Exploring the influence of the genetic region on *HMGCR* expression level associations with CHD, left hand side selecting from ± the genetic region, right hand side selecting from either the upstream region or the downstream region; colours represent z-statistics and stars indicate significant associations at a type 1 error rate of 0.05

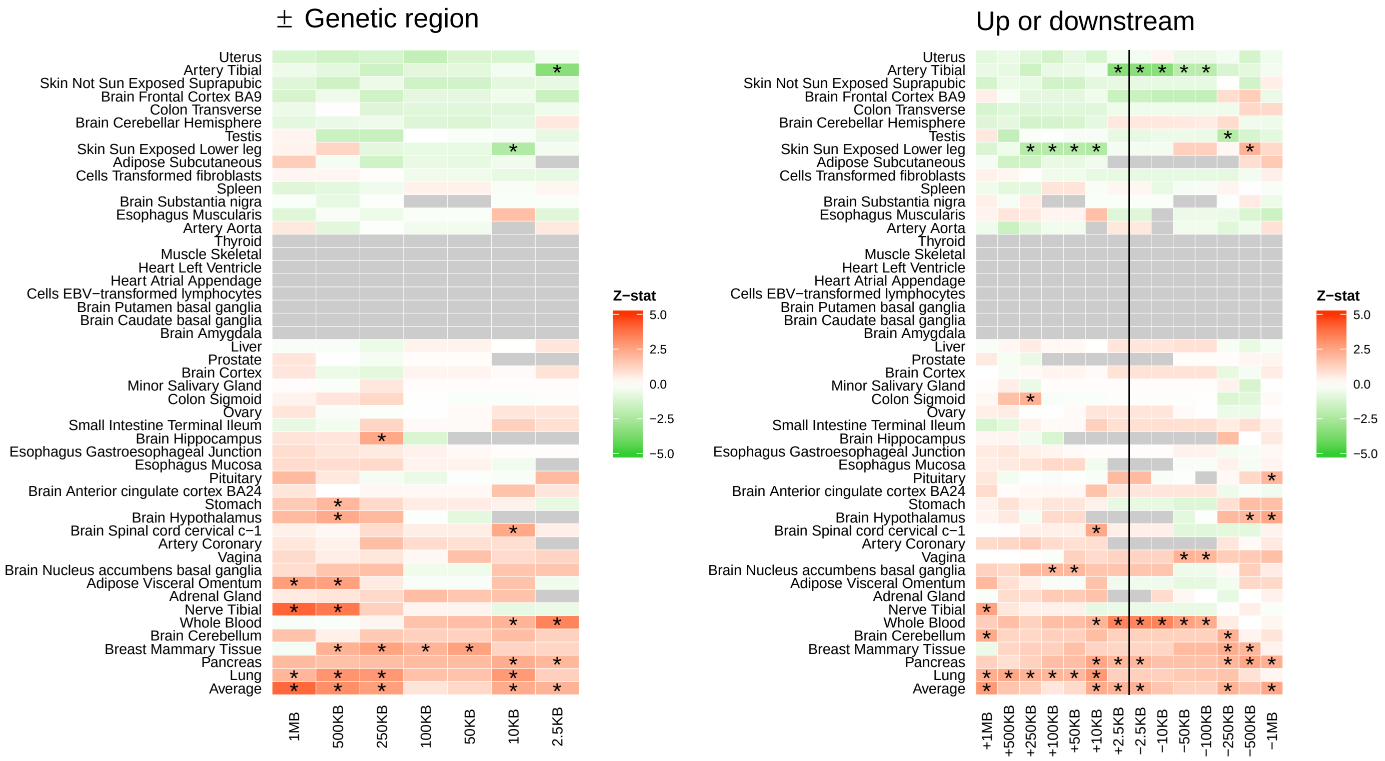


**Appendix Figure 13:** Exploring the influence of the genetic region on *PCSK9* expression level associations with CHD, left hand side selecting from ± the genetic region, right hand side selecting from either the upstream region or the downstream region; colours represent z-statistics and stars indicate significant associations at a type 1 error rate of 0.05

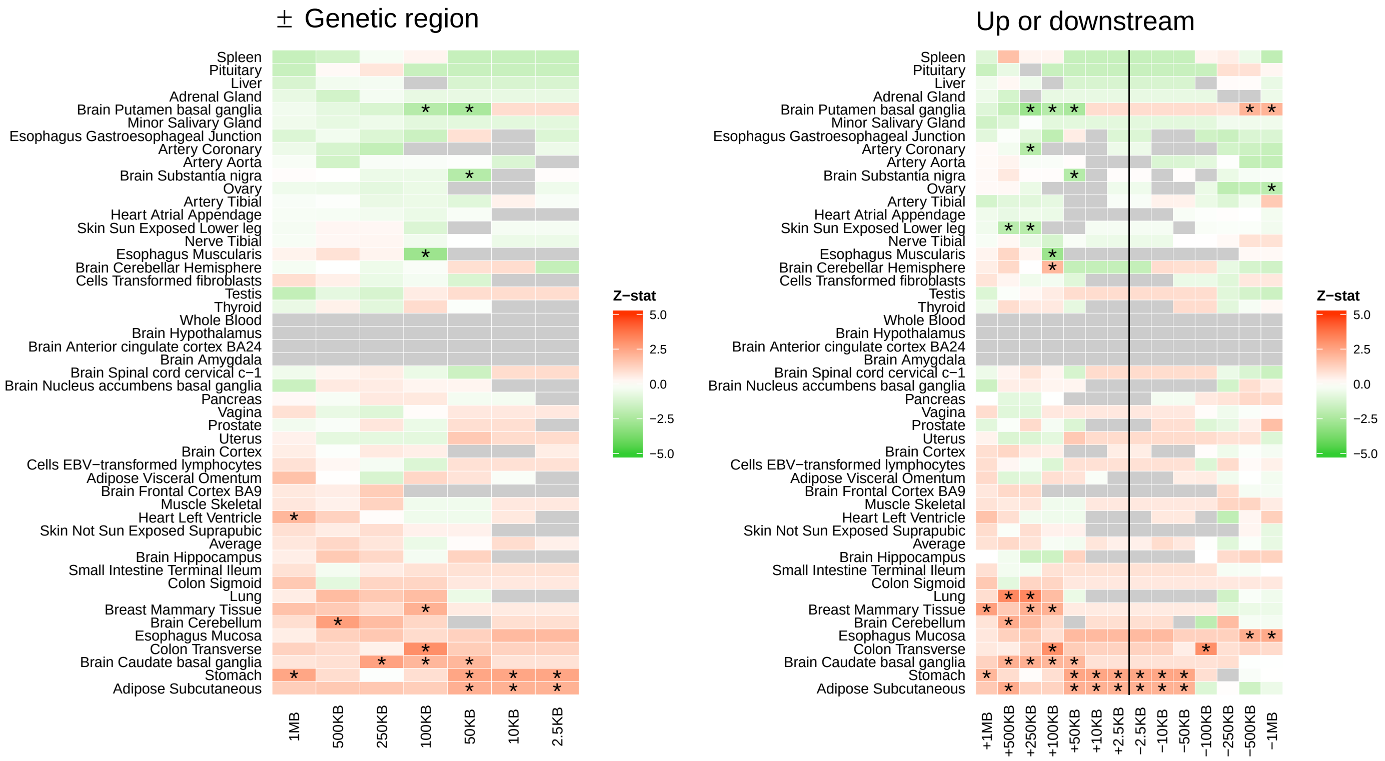


**Appendix Figure 14:** Exploring the influence of the genetic region on *NPC1L1* expression level associations with CHD, left hand side selecting from ± the genetic region, right hand side selecting from either the upstream region or the downstream region; colours represent z-statistics and stars indicate significant associations at a type 1 error rate of 0.05

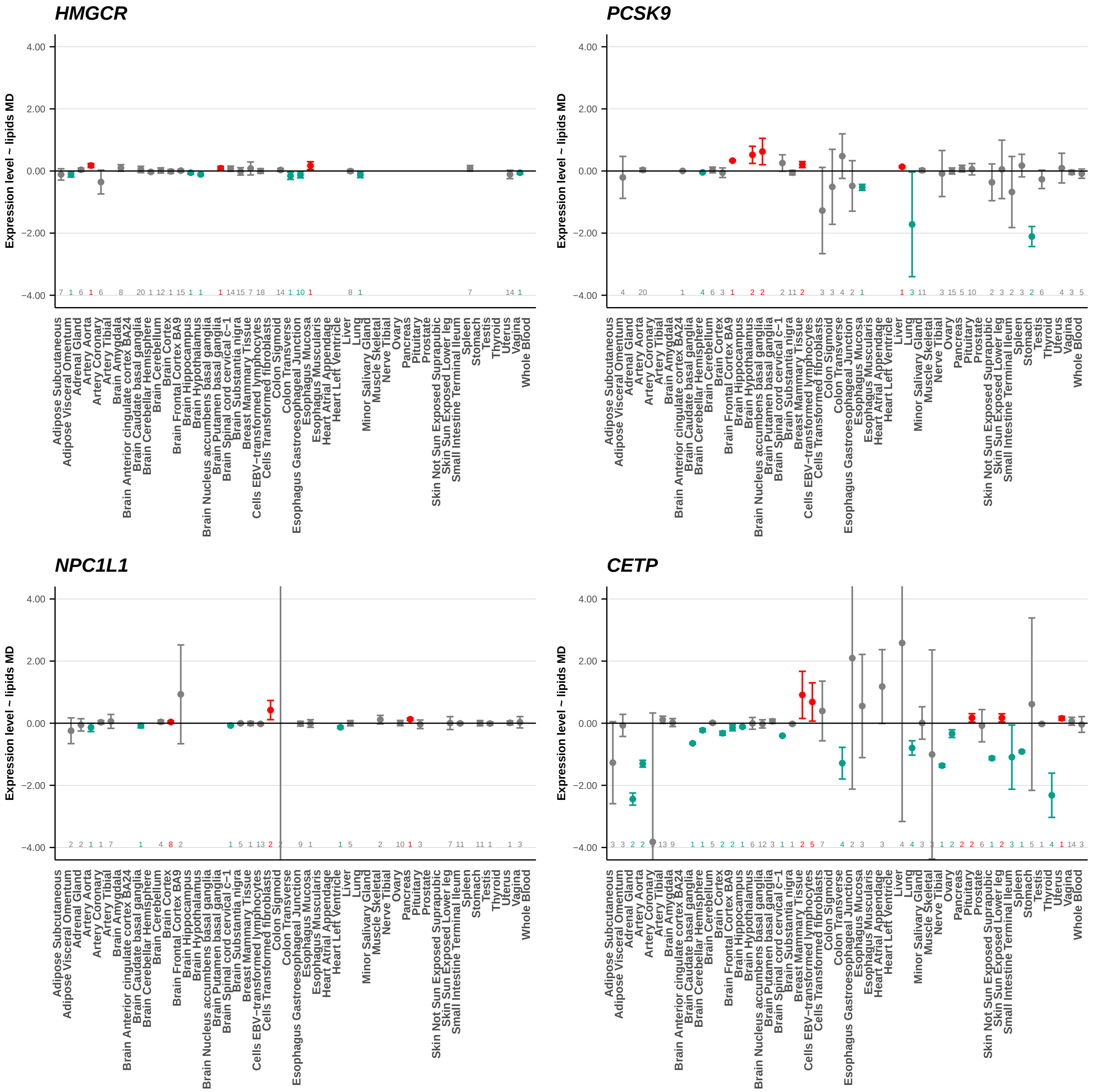


**Appendix Figure 15:** Mendelian randomization estimates of the expression level effects on lipids. Instruments were taken from the *HMGCR* locus (top left), the *PCSK9* locus (top right panel), *NPC1L1* locus (bottom left), and *CETP* locus (bottom right). eQTL data were available from GTEx [7] and lipids from GLGC [2]; multiplicative random-effects standard error estimates were corrected for LD using the "EUR" 1000 genomes panel [8] and the estimator proposed in [9, 10] with an Egger analytical correction for pleiotropy .

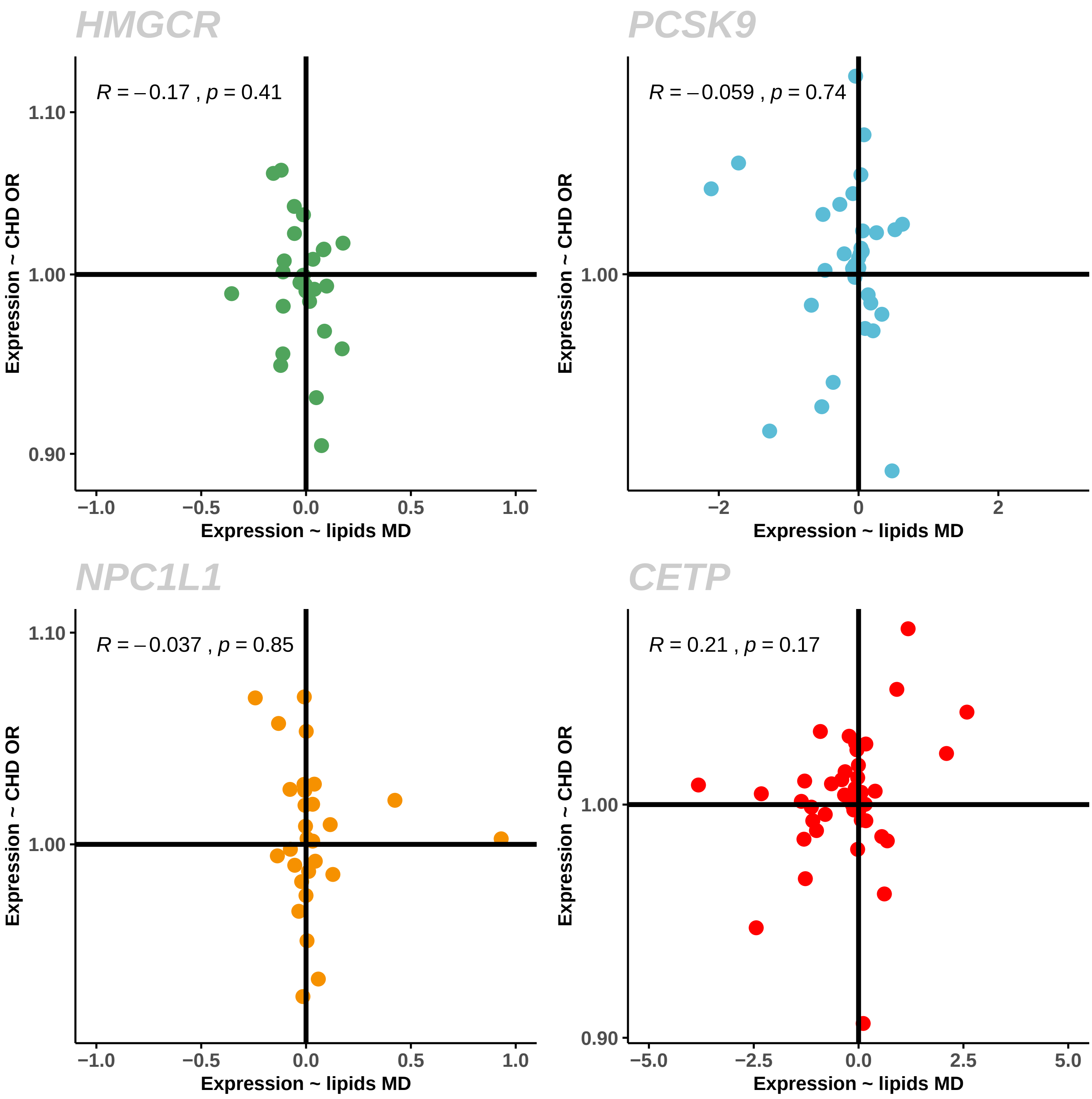


**Appendix Figure 16:** Exploring concordance between expression level MR estimates with lipids and with CHD (after Egger correction).

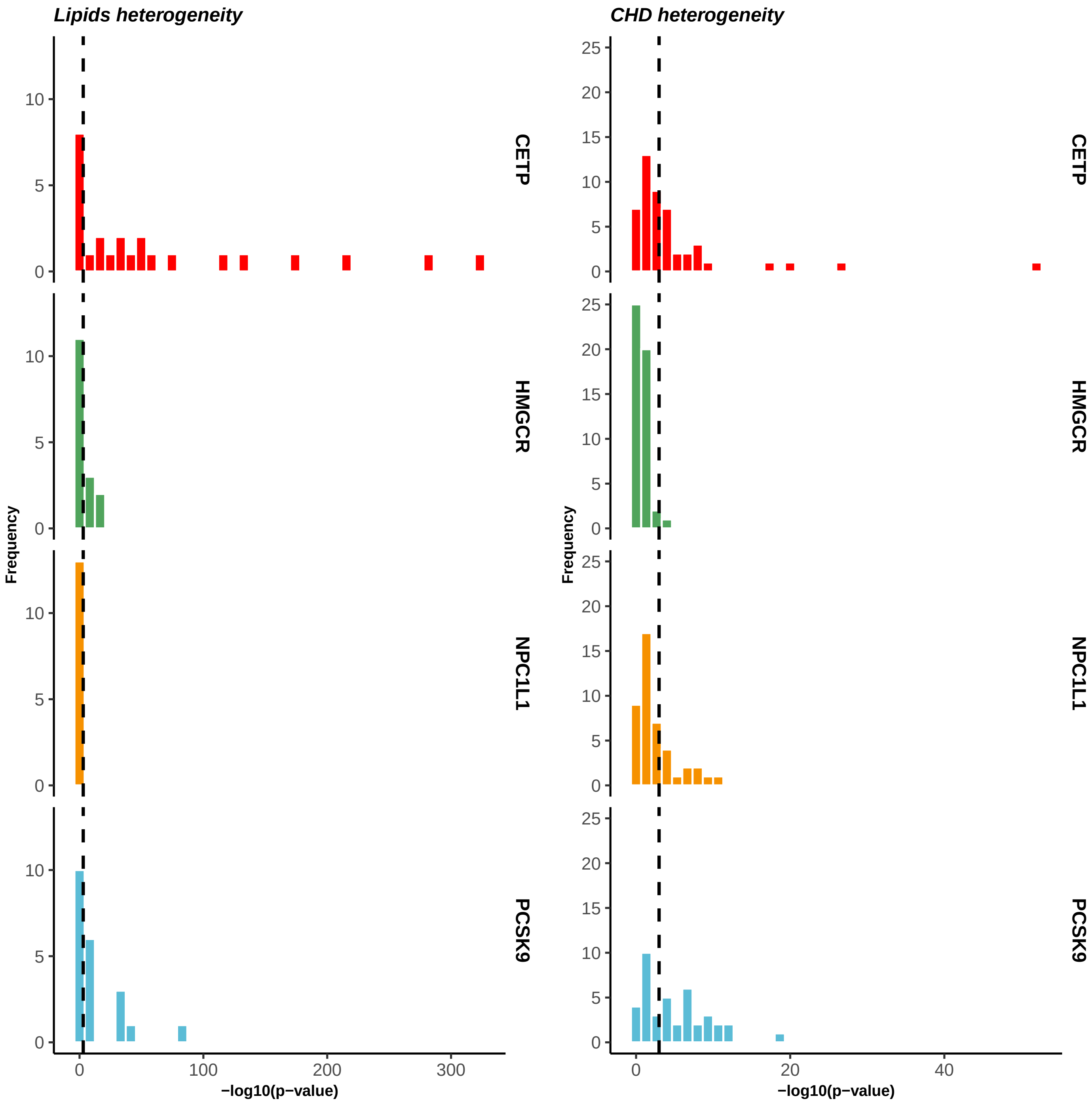


**Appendix Figure 17:** Heterogeneity p-values (Q-statistics) of MR-egger adjusted estimates of the tissue specific expression level effects on lipids and CHD. Vertical line indicates a p-value of 0.0010 taking as a conservative indicator of heterogeneity.

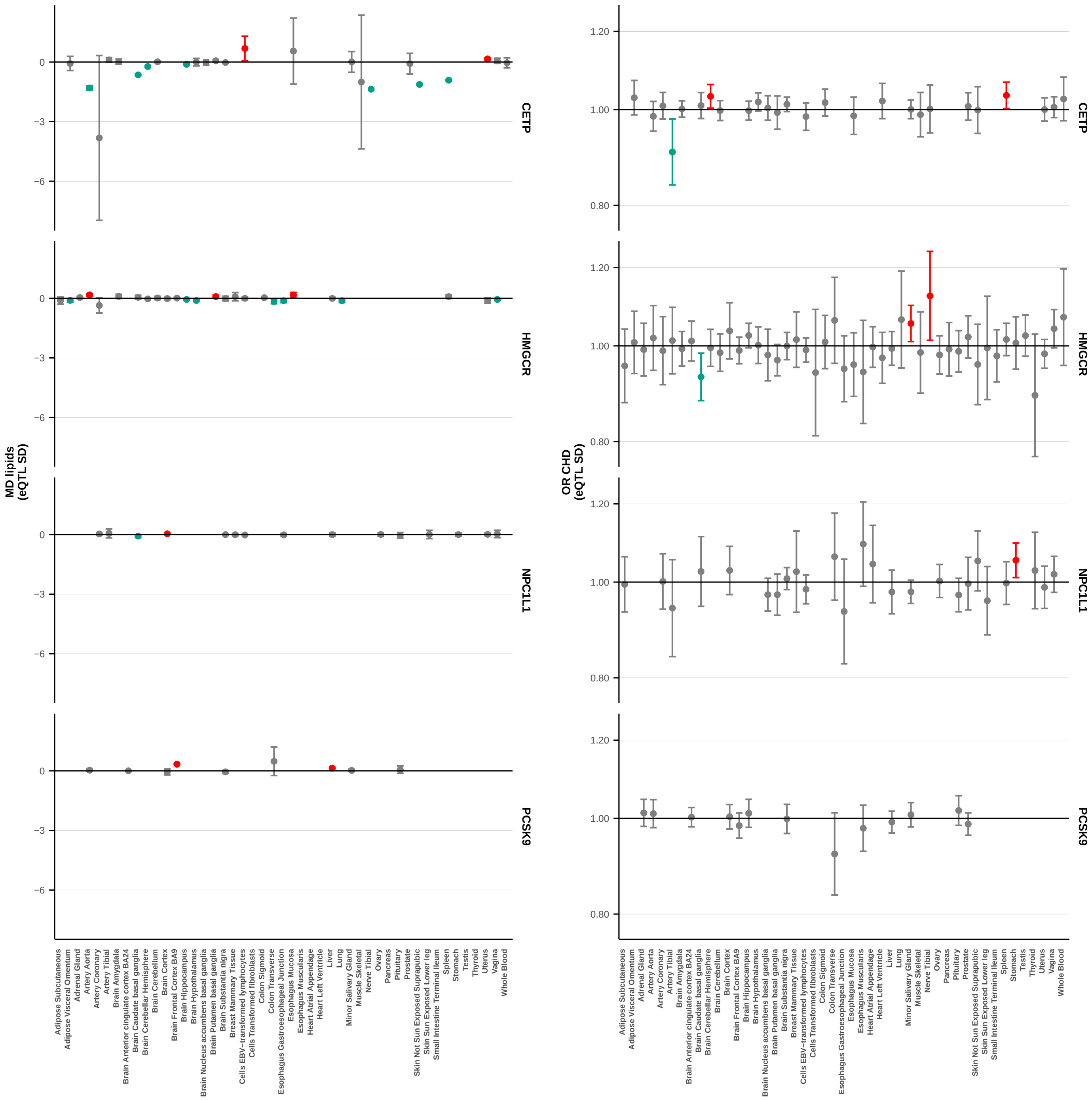


**Appendix Figure 18:** Expression level effects on lipids and CHD, after excluding heterogeneous tissues

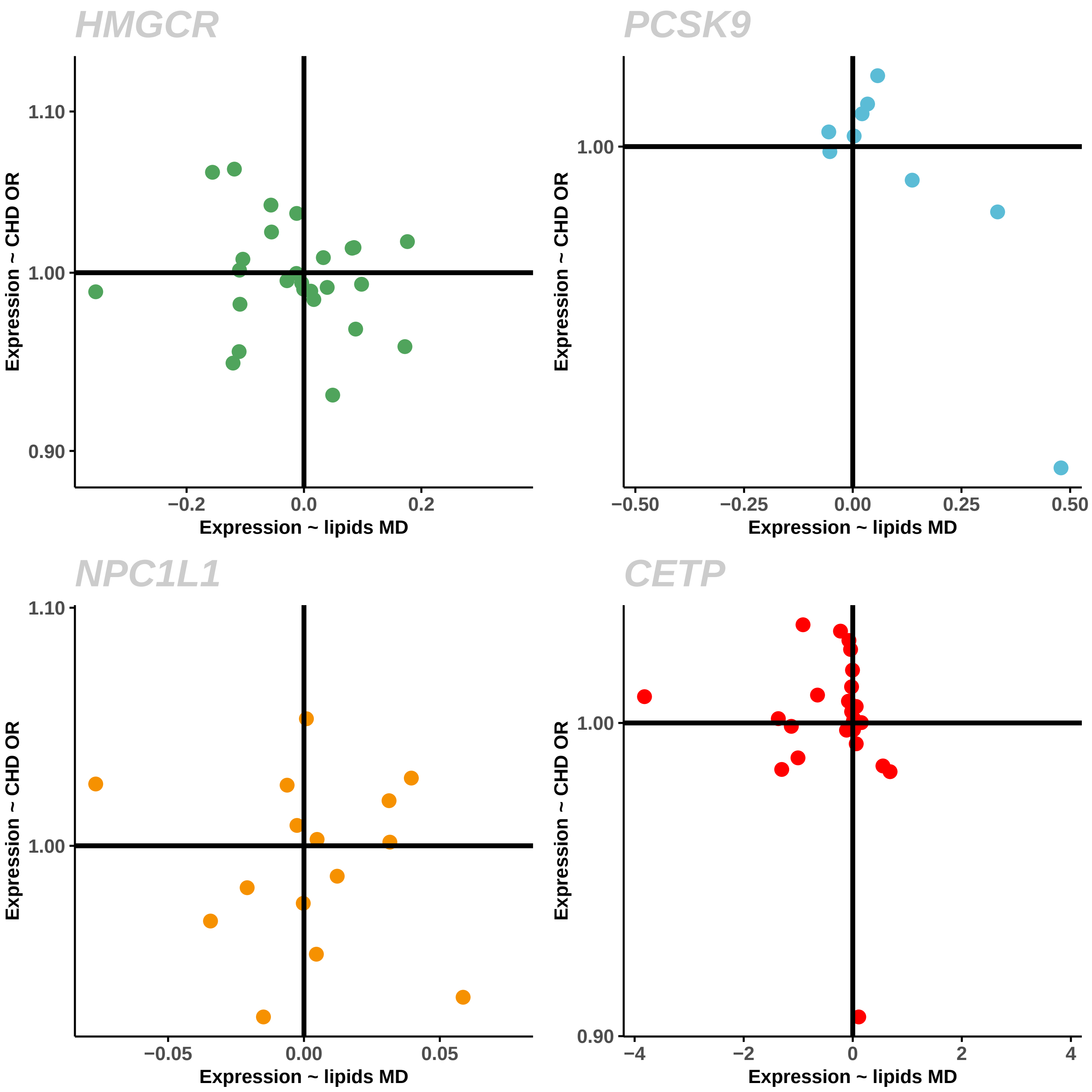


**Appendix Figure 19:** Expression effects on lipids and CHD, after excluding heterogeneous tissues (concordance)

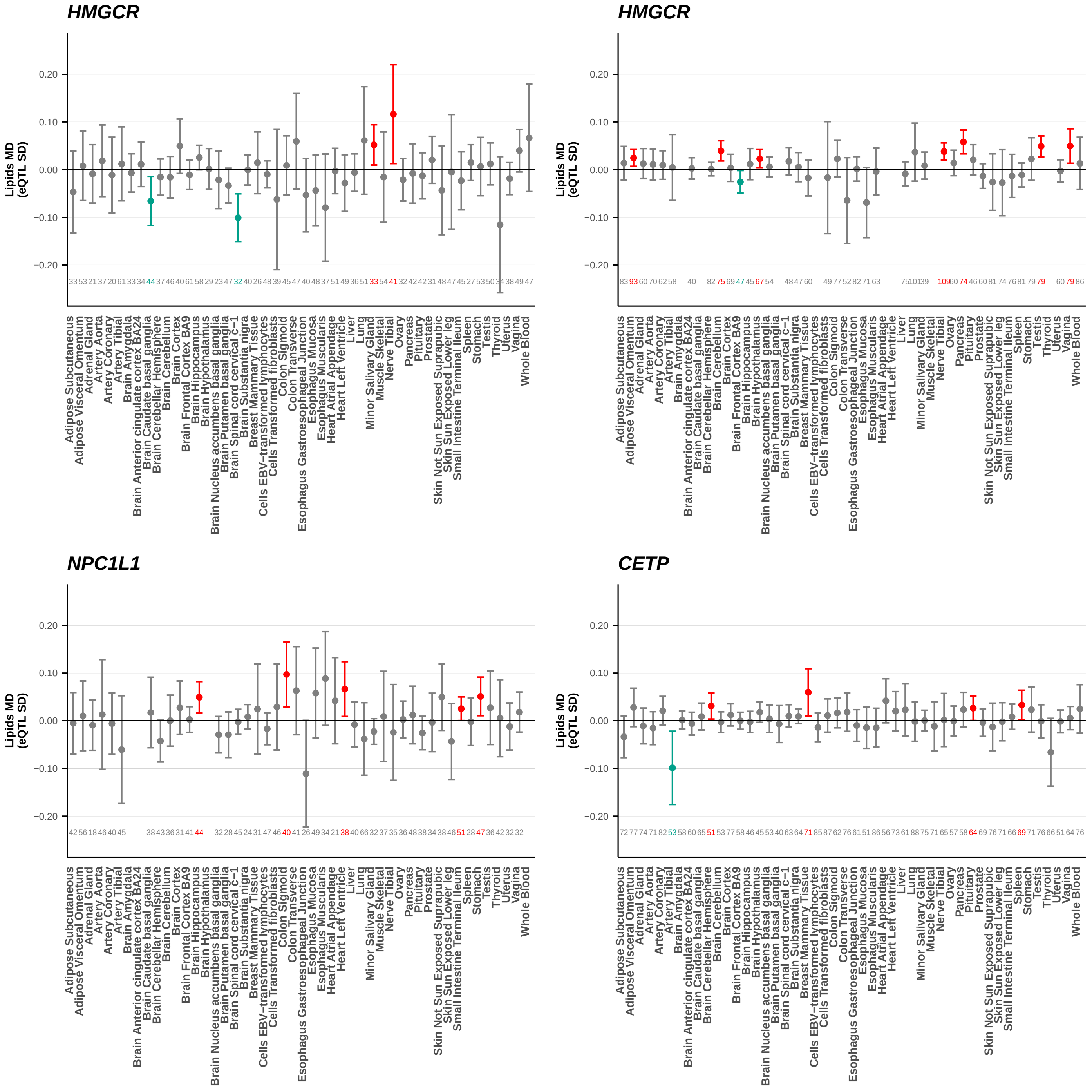


**Appendix Figure 20:** Expression lipid effects after removing heterogeneous variants

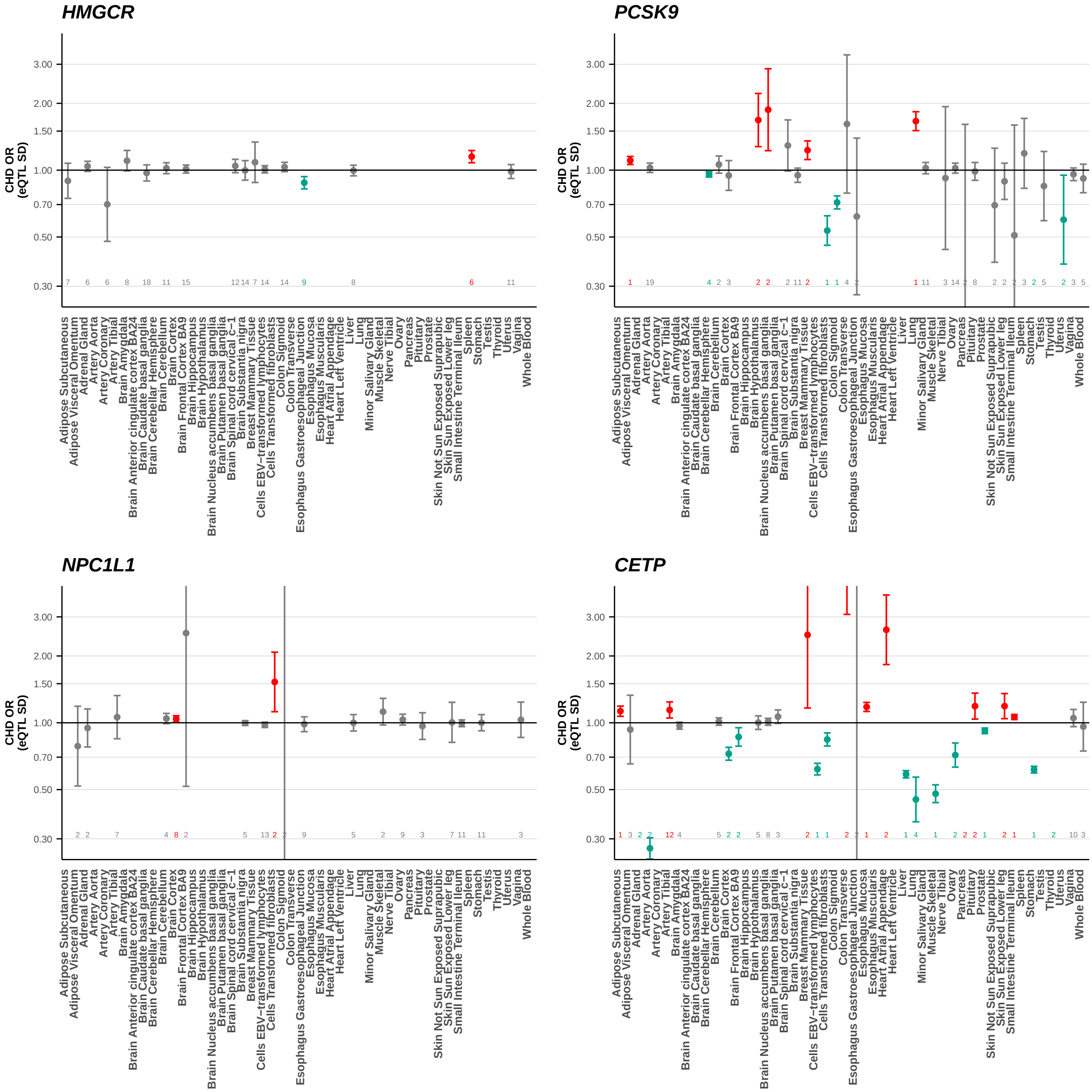


**Appendix Figure 21:** Expression level CHD effects after removing heterogeneous variants

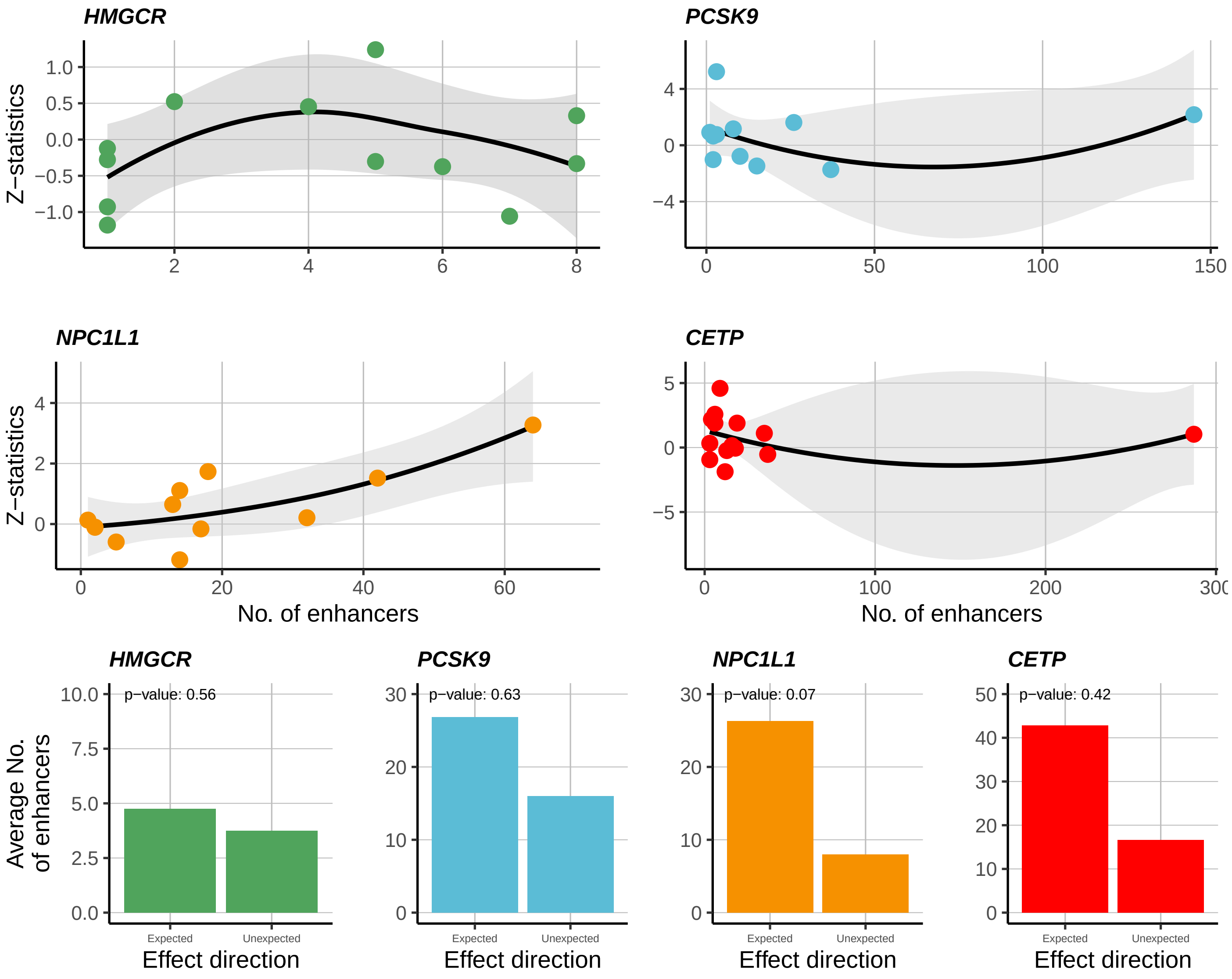


**Appendix Figure 22:** The influence of enhancer variants on the expression level association with CHD

### References


[1] Benjamin B Sun et al. “Genomic atlas of the human plasma proteome.” In: *Nature* 558 (7708 June 2018), pp. 73–79. ISSN: 1476-4687. DOI: 10.1038/s41586-018-0175-2.

[2] Cristen J Willer et al. “Discovery and refinement of loci associated with lipid levels.” In: *Nature genetics* 45 (11 Nov. 2013), pp. 1274–1283. ISSN: 1546-1718. DOI: 10.1038/ng.2797.

[3] Majid Nikpay et al. “A comprehensive 1,000 Genomes-based genome-wide association meta-analysis of coronary artery disease.” In: *Nature genetics* 47 (10 Oct. 2015), pp. 1121–1130. ISSN: 1546-1718. DOI: 10.1038/ng.3396.

[4] Heribert Schunkert et al. “Large-scale association analysis identifies 13 new susceptibility loci for coronary artery disease.” In: *Nature genetics* 43 (4 Mar. 2011), pp. 333–338. ISSN: 1546-1718. DOI: 10.1038/ng.784.

[5] Karsten Suhre et al. “Connecting genetic risk to disease end points through the human blood plasma proteome.” In: *Nature communications* 8 (Feb. 2017), p. 14357. ISSN: 2041-1723. DOI: 10.1038/ncomms14357.

[6] Lisanne L Blauw et al. “CETP (Cholesteryl Ester Transfer Protein) Concentration: A Genome-Wide Association Study Followed by Mendelian Randomization on Coronary Artery Disease.” In: *Circulation. Genomic and precision medicine* 11 (5 May 2018), e002034. ISSN: 2574-8300. DOI: 10.1161/CIRCGEN.117.002034.

[7] GTEx Consortium. “Genetic effects on gene expression across human tissues.” In: *Nature* 550 (7675 Oct. 2017), pp. 204–213. ISSN: 1476-4687. DOI: 10.1038/nature24277.

[8] The 1000 Genomes Project Consortium. “A global reference for human genetic variation.” In: *Nature* 526 (7571 Oct. 2015), pp. 68–74. ISSN: 1476-4687. DOI: 10.1038/nature15393.

[9] Stephen Burgess, Frank Dudbridge, and Simon G Thompson. “Combining information on multiple instrumental variables in Mendelian randomization: comparison of allele score and summarized data methods.” In: *Statistics in medicine* 35 (11 May 2016), pp. 1880–1906. ISSN: 1097-0258. DOI: 10.1002/sim.6835.

[10] Stephen Burgess, Verena Zuber, Elsa Valdes-Marquez, Benjamin B Sun, and Jemma C Hopewell. “Mendelian randomization with fine-mapped genetic data: Choosing from large numbers of correlated instrumental variables.” In: *Genetic epidemiology* 41 (8 Dec. 2017), pp. 714–725. ISSN: 1098-2272. DOI: 10.1002/gepi.22077.
